# The Friendship Paradox across animal social systems is governed by network structure and biological features

**DOI:** 10.64898/2026.03.24.713537

**Authors:** Eloise F. Newman, Sarah C.L. Knowles, Josh A. Firth

## Abstract

A population’s social structure, often represented as a social network, shapes fundamental biological processes including the spread of disease, information, and behaviour. The ‘Friendship Paradox’ is a network phenomenon whereby the average individual has fewer ‘friends’ than their ‘friends’ do. This effect can be quantified as relationship disparity (the difference between an individual’s connectedness and those they are connected to) which captures the local social environment. Previous work has shown that such relationship disparity can be exploited in effective outbreak monitoring, targeted health interventions and optimized contact tracing. Yet, how its magnitude varies across real-world social networks remains poorly understood. Here, we analyse relationship disparity across 391 empirical animal social networks to test how intrinsic network properties and biological attributes predict its extent. We find that smaller and sparser networks exhibit stronger relationship disparity, and it is further shaped by both species-specific differences and shared phylogenetic history. After controlling for variation in individual sociability, relationship disparity was more strongly accentuated in mammalian direct physical contact networks than indirect social association networks, whilst the corresponding finding was not detected among Sauropsids. Phylogenetic and species identity largely explained variation in relationship disparity indicating biological drivers. Together, these findings demonstrate that both network structure and biological attributes shape relationship disparity in animal social systems, providing a foundation for predicting how higher-order network architecture influences social processes such as contagion.

**Significance Statement:** In natural populations, social connections are unevenly distributed, often resulting in a small subset of individuals that are highly connected while many are relatively peripheral. The ‘Friendship Paradox’ is a measure of relationship disparity between individuals and their local social environment. Understanding how features of the social network and biological system are associated with relationship disparity can contribute to understanding what shapes social behaviour. Relationship disparity may not just be an emergent network property but could reflect a higher level of social structuring, and therefore shape processes that depend on social contacts. Our findings demonstrate the value of comparative network analysis for generating insights into fundamental principles structuring real-world societies.

## Introduction

Animal populations are structured by patterns of social interaction (1). Representing these patterns of interaction as social networks provides a powerful way to explore how they shape the evolutionary and ecological drivers and consequences of sociality (1, 2). One phenomenon that emerges from simple patterns of interaction within social networks is Feld’s Friendship Paradox, which posits that the average individual has fewer ‘friends’ (social connections) than their friends do (figure 1). This paradox arises because real-world social networks are not fully connected, and because highly connected individuals, although relatively rare, are overrepresented among any given individual’s social contacts (3–5). The ‘Friendship Paradox’ can be quantified as relationship disparity: the extent to which an individual’s social partners have more (or fewer) relationships than the individual itself, calculated at the individual level and averaged over the whole network (4, 6). Relationship disparity is therefore a parameter that captures the extent to which an average individual’s connections are better connected than they are, or the strength of the friendship paradox in a social network (figure 1). Although theory predicts that social network structure should influence the strength of the Friendship Paradox (4), it remains unclear how much relationship disparity is observed in natural animal social networks, whether it varies systematically among taxa, interaction types, or social contexts, and what consequences such variation might have for social processes.

**Figure 1.**
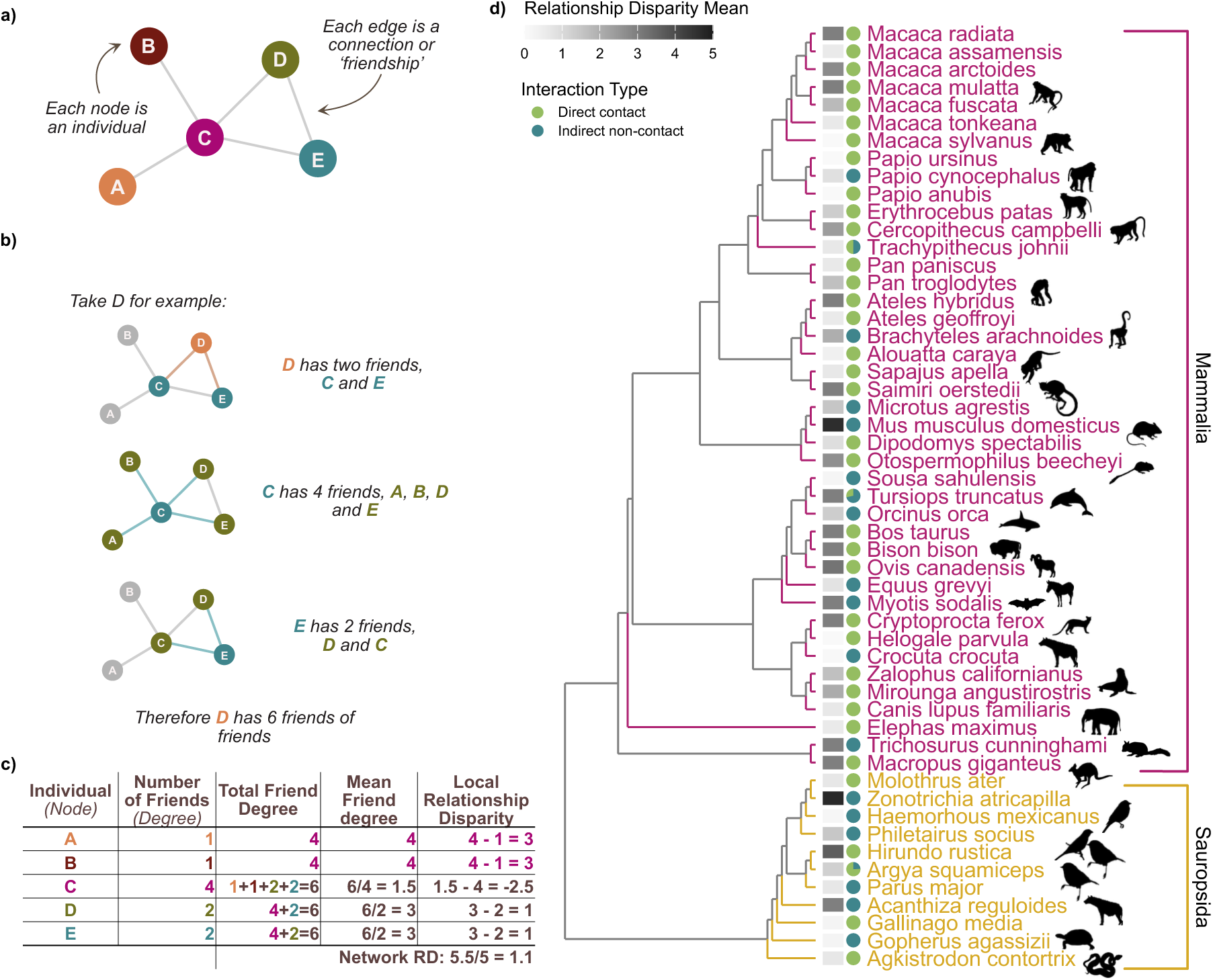
Calculating relationship disparity on a network and phylogeny of real-world animal social networks. a) Structure of an example network; b) each node is taken in turn, and their connections (friends) and connections’ connections (friends of friends) are calculated which is shown in c) Local relationship disparity is the mean connectedness of a node’s direct social partners minus the connectedness of the focal node. Network-level relationship disparity is the mean of these local values across all nodes. For unweighted networks, connectedness is measured as degree; for weighted networks, it is measured as node strength, calculated as the sum of a node’s edge weights. d) Phylogeny of the taxonomic distribution of animal social networks and relationship disparity. Species binomial names are listed at the tip of the linear tree. The greyscale heatmap represents the average relationship disparity across all the network(s) of the respective species. Mammalia (“pink”) and Sauropsida (“yellow”) are depicted by colour of species name. Animal illustrations are from Phylopic in the closest taxonomic relation if unable to find exact taxa with specific silhouette images citations listed in the supplementary.

Quantifying relationship disparity in real-world social networks provides insight into the forces shaping social structure, as systematic differences between individuals and their social partners in connectedness can arise from variation in sociability, dominance, movement, or ecological constraints (7), and in turn influence patterns of interaction, competition, and cooperation within groups (8–11). Relationship disparity also has important consequences for how disease, information, and behavioural change spread, as its magnitude determines how effectively the Friendship Paradox can be exploited for surveillance and intervention (3, 4, 12–17). In humans, for example, an application of the friendship paradox concept (13) showed that recruiting the social partners of randomly selected individuals allowed peaks in influenza prevalence to be detected nearly three weeks earlier than under random sampling alone (13), and subsequent work has shown that individuals with the highest network centrality can act as particularly sensitive ‘sensors’ of outbreaks (14, 15, 18, 19).

In practice, however, identifying such highly central individuals is often infeasible because it requires detailed knowledge of the entire network. Node-level measures of relationship disparity offer a practical alternative as they can be calculated for any individual using only local information on their social partners. Measures based on relationship disparity have already been used to detect influenza outbreaks from social media (20), identify early COVID-19 infections from phone data (15), track shifts in voting intentions (21), and monitor adoption of health behaviours (22–24). The effectiveness of these approaches depends on the underlying network structure: in networks with strong relationship disparity, sampling a small number of well-connected individuals can both improve surveillance and even allow the estimation of broader network properties (3, 18). Yet we still lack a clear understanding of how strong relationship disparity is in real-world networks and why it varies among social systems and contexts.

Comparative studies across diverse animal social networks provide a powerful way to address this gap (25, 26). The Animal Social Network Repository (ASNR) contains over 1100 animal social networks spanning diverse taxa and interaction types (27, 28), enabling analysis of the fundamental drivers of relationship disparity, and a better understanding of when Friendship Paradox based strategies will be most effective in outbreak monitoring, promoting or tracking social learning, and contact tracing in both human and non-human populations.

Here, we use a large comparative empirical dataset of real animal social networks to examine how the Friendship Paradox varies across social systems and what network and biological traits predict this variation. We further test whether the observed level of relationship disparity in each network exceeds or falls below expectations based on individual sociability alone (i.e. degree distribution), revealing taxonomic and interaction-type features associated with accentuated or weakened relationship disparity. Collectively, our results shed light on the drivers of variation in animal social structure and indicate when Friendship Paradox-based strategies might be most effectively applied in real-world settings.

## Results

We compiled social networks of wild vertebrate populations from the ASNR containing 10 to 1100 individuals (excluding networks with >50% isolated nodes). Social interaction types were aggregated into direct physical interactions (e.g. mating, grooming) or indirect interactions (e.g. co-occurrence, group associations) (see supplementary text). The final dataset consisted of 391 social networks of 53 species, the majority of which derived from mammals (n=42, networks = 283) and non-mammal vertebrates Sauropsida (n=11, networks = 108), composed of avian (n=9*, networks = 62) and non-avian reptiles (n=2, networks = 46) (Fig. 1).

We began by measuring relationship disparity across networks, and then asked which network and biological features best explain variation among social systems. Relationship disparity was calculated for each individual as the difference between the mean number of connections of its social partners and its own number of connections; network-level relationship disparity was defined as the mean of this quantity across all individuals and can be interpreted as the average experience of relationship disparity in a network (figure 1, supplementary text). This is a separate measure from degree assortativity, or degree heterogeneity (see Supplementary Information)

### Network structural properties predict relationship disparity

As all network-level relationship disparity values were positive and skewed, they were natural log-transformed prior to analysis to better approximate a normally distributed response variable. Parameter estimates are therefore reported on the log scale, whereas figures show back-transformed values. We first examined two basic network descriptors, (i) network size and (ii) whether networks were weighted or unweighted, as predictors of relationship disparity in a phylogenetic model with phylogeny and species identity as a random effect. This revealed that relationship disparity tended to be weaker in larger networks (Est=-0.08 +/- 0.08, 95% credible interval, CI: -0.23 to 0.07), although this effect was uncertain. Weighted networks showed no difference in relationship disparity than unweighted networks (weighted: Est=0.03 +/- 0.56, CI: -1.06 to 1.12). Both network size and weighting were retained as control variables in subsequent models.

We then investigated how finer-scale network structure predicted relationship disparity. When considered individually, mean degree (the average number of connections per individual) was positively associated with relationship disparity (Est=0.37 +/- 0.12, CI: 0.13 to 0.62), while edge density (the proportion of possible connections that were realised) showed a strong negative association with relationship disparity (Est=-0.57 +/- 0.12, CI: -0.80 to -0.33). Group cohesion, a measure of how many links must be removed to divide the network into two components, showed no clear relationship with relationship disparity when modelled individually (Est=-0.031 +/- 0.11, CI: -0.24 to 0.18). In a joint model including mean degree, edge density, group cohesion and the basic network descriptors (size and weighting), all three structural predictors were strongly associated with relationship disparity: networks with higher mean degree (Est=0.44 +/- 0.13, CI: 0.19 to 0.71), lower edge density (Est=-0.88 +/- 0.15, CI: -1.19 to -0.58), and greater group cohesion (Est=0.36 +/- 0.13, CI: 0.09 to 0.63) had higher relationship disparity. In summary, after accounting for network size and weighting, relationship disparity was generally higher in sparser social networks with many connections per individual and more cohesive subgroup structure (figure 2).

**Figure 2.**
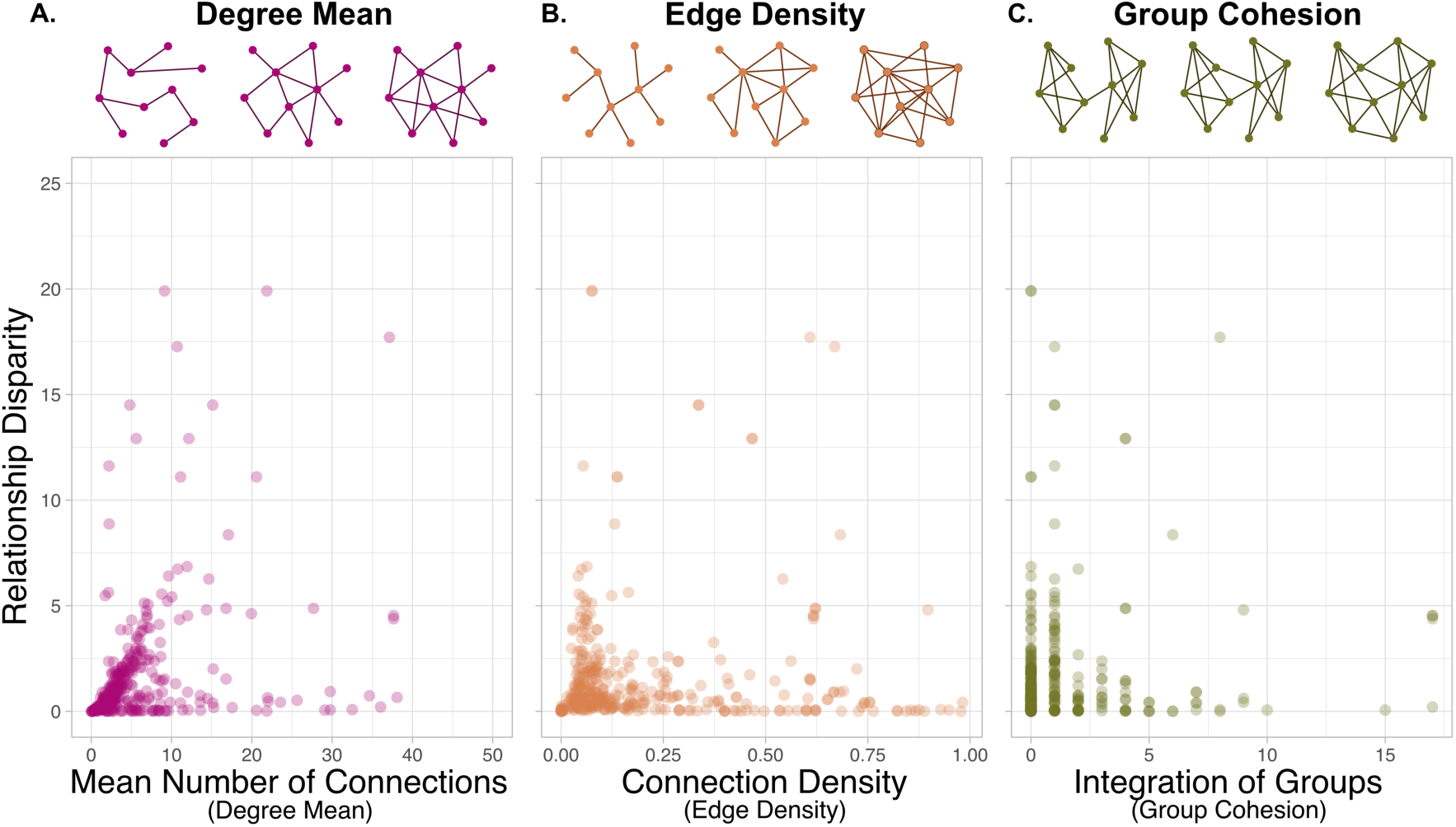
The degree of relationship disparity as a function of network features. a) Degree mean, b) edge density and c) group cohesion. Illustrations of changes in the network are presented above each plot, with raw data plotted below.

### Phylogeny and species identity, but not interaction type, predicts relationship disparity

To test whether biological attributes predict variation in relationship disparity beyond network structure, we fitted a phylogenetic mixed-effects model with relationship disparity as the response and interaction type as a predictor. Phylogenetic non-independence among species was incorporated through a covariance matrix, species was included as a random effect, and network size, weighting, mean degree, edge density, and group cohesion were included as covariates. Social networks based on direct social interactions showed slightly weaker relationship disparity than networks based on indirect social interactions, though this difference was not statistically significant (figure 3a; Est=-0.33±0.37, 95% CI: -1.05 to 0.38). Relationship disparity varied substantially among species (species SD=1.85±0.30, 95% CI: 1.28 to 2.49) and with shared phylogenetic ancestry (phylogeny SD=0.69±0.56, 95% CI: 0.02 to 2.04). Posterior variance decomposition indicated that species-specific effects explained the largest proportion of the observed variation in relationship disparity (60.3%), followed by within-species residual variance (28.0%) and phylogenetic effects (11.7%) (supplementary figure S4 and supplementary table S41).

**Figure 3.**
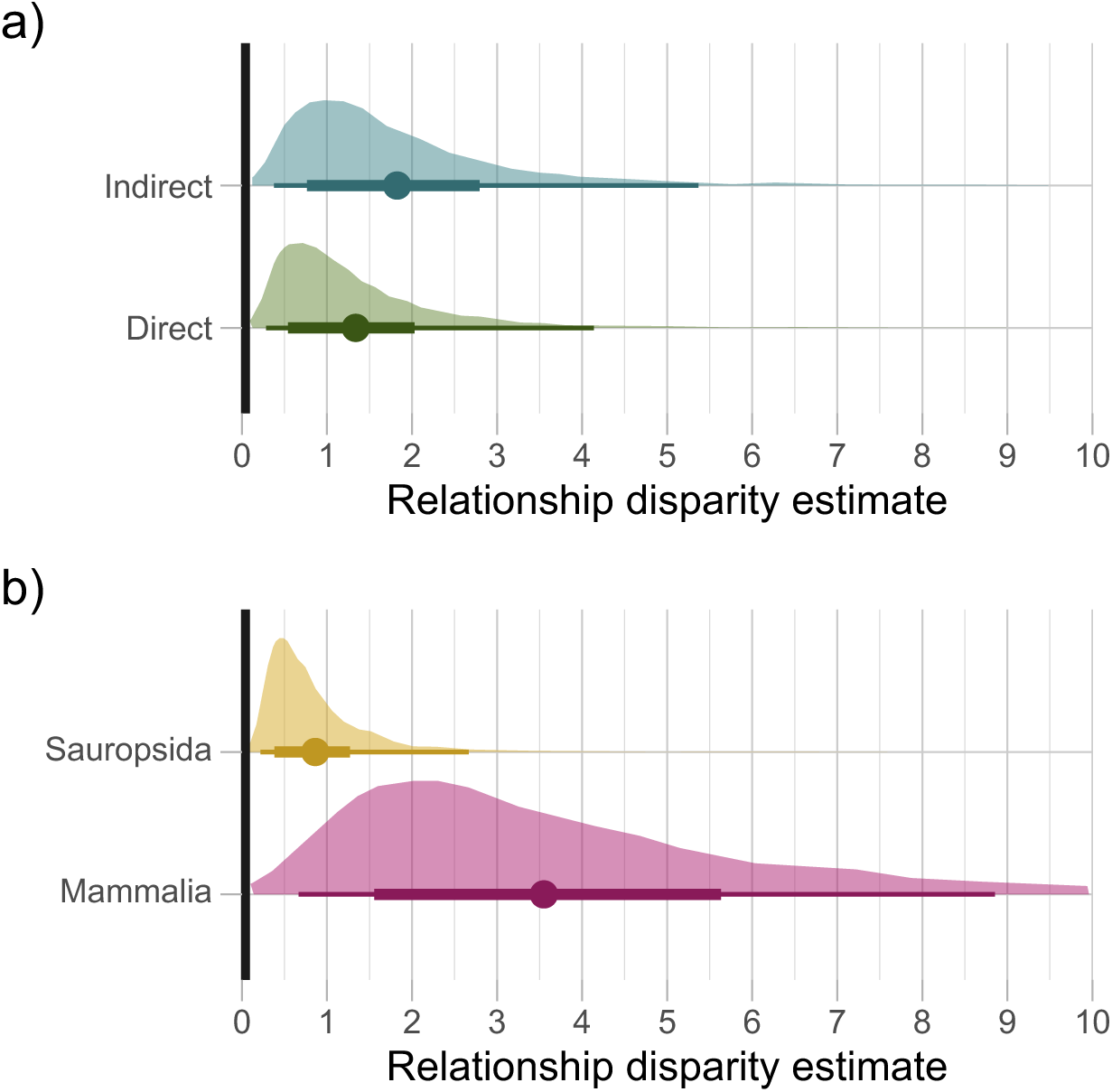
Posterior parameter estimates of relationship disparity according to biological network properties: a) interaction type of direct physical networks and indirect non-contact networks; and b) two major taxonomic groupings of Mammalia and Sauropsida from two taxonomic specific models. Relationship disparity values were natural log-transformed for analysis, and network features centred and scaled for modelling. Results were extracted from models as posterior parameter draws as reported and are back transformed to original scale for clarity and interpretation. Points represent the posterior mean estimate of relationship disparity for each network type, and bars represent 66% (around one standard deviation) and 95% (around two standard deviations) credible intervals. Distributions show the full posterior uncertainty in the estimated mean relationship disparity for each group.

### Relationship disparity deviates from expectations based on individual variation in sociability

Observed relationship disparity values could in principle arise solely from network structural properties, particularly individual variation in sociability (degree distributions) or could instead reflect additional structure in how individuals connect to specific other individuals. To test whether relationship disparity values differed in magnitude more than expected based solely on individual variation in number of connections, and whether biological attributes could predict this magnitude, we constructed permutation-based null models for each network, randomly rewiring edges 1,000 times while holding the degree distribution, network size and edge density constant (and preventing self-loops or multiple edges). For each network, we calculated a z-score for relationship disparity as the deviation of the observed value from the null mean measured in units of the null standard deviation (supplementary figure S3). If the observed value is significantly different from the null distribution of values in a z-test, the network’s relationship disparity value is considered accentuated – either significantly greater or less than what is expected. With a 95% credible level, 5% of permutation tests are expected to be significant due to chance (type 1 error). However, 47.8% of all social networks’ real relationship disparity values were more extreme (either significantly stronger (9.7%), or weaker (38.1%)) than the null expectation based on their degree distribution, network size and edge density.

### Direct and indirect social networks do not deviate in expected relationship disparity

The extent to which social networks differed from the relationship disparity expected from their own degree distributions showed no clear overall difference between networks derived from direct and indirect social interactions in estimated mean z-score (Est=0.57±0.58, 95% CI: -0.56 to 1.73;, figure 4a). Differences were primarily attributed to specific species (species SD=7.03±0.70, 95% CI: 5.77 to 8.50) rather than a smaller significant impact of shared ancestry (phylogeny SD=0.75±0.63, 95% CI: 0.02 to 2.36). This indicates that specific species-level factors (such as ecology or behaviour) may drive the observed value which have not been accounted for specifically in the model. The model explained an estimated 70.3% of variation in the response (Bayes R2 = 0.703±0.016, 95% CI: 0.67 to 0.73). In posterior variance decomposition, species identity accounted for 90.5% of the total variance, with 7.7% attributed to within-species residual variance effects, whilst phylogeny was 1.8%. To see if differences between the two clades alone could explain these values, a model without phylogeny but species identity was run (supplementary tables S20-32). No significant difference was found between sauropsidan and mammalian networks (Est=-0.2542±0.91, 95% CI: -2.01 to 1.56) in non-phylogenetically controlled models with clade included as a categorical variable (supplementary table S32).

**Figure 4.**
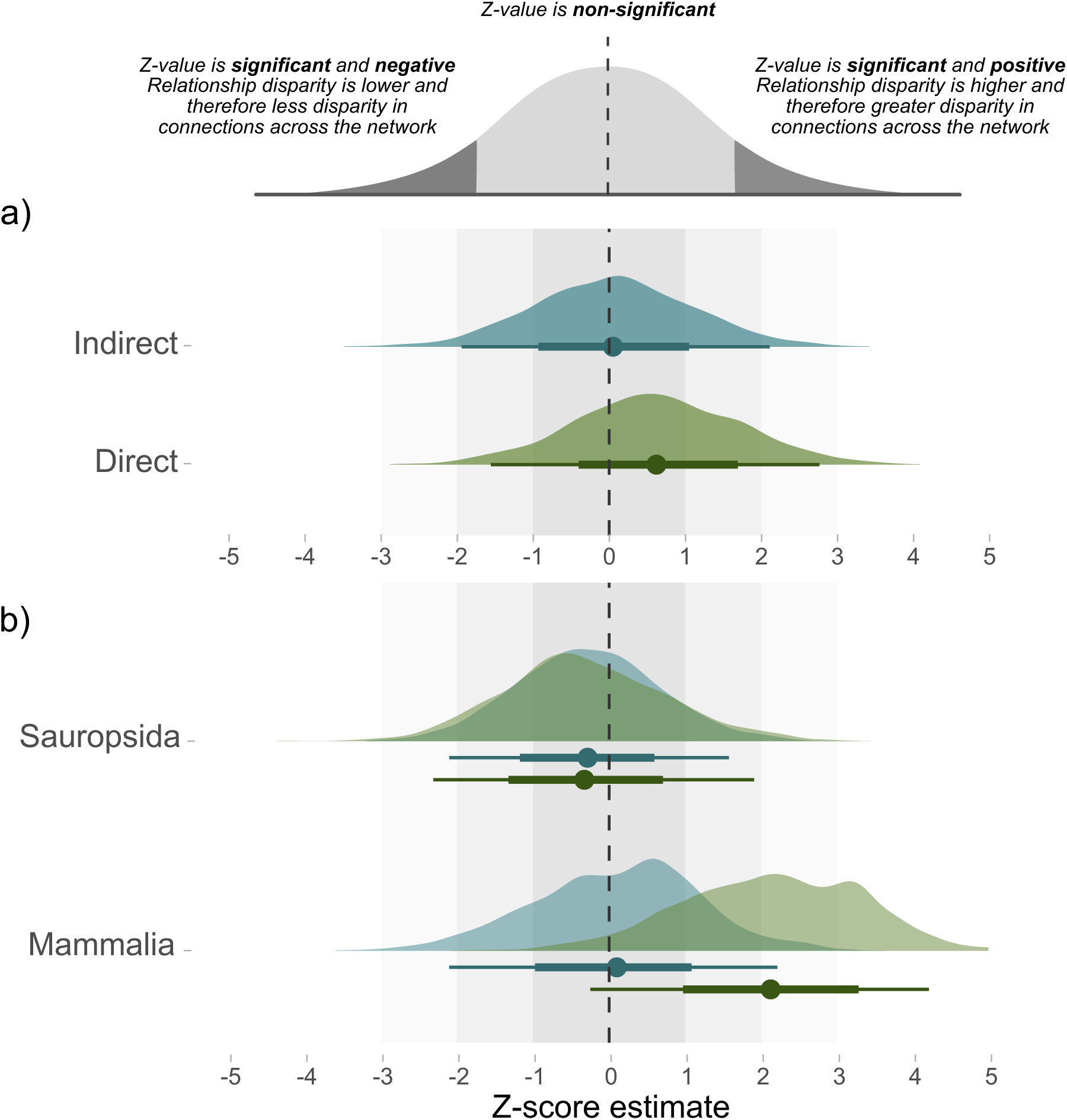
Posterior parameter estimates of relationship disparity accentuation split into direct and indirect interaction networks. a) Parameter estimates are from the overall phylogenetic model considering interaction type and species identity as a random effect; and b) shows the z-score parameter estimates for direct (green) and indirect (blue) interaction networks from two separate models of containing only Sauropsida and Mammalia respectively. Mammalian direct and indirect networks are significantly different. Points represent the posterior estimated mean z-score for each network type, and bars represent 66% (one standard deviation) and 95% (two standard deviations) credible intervals. Distributions show the full posterior uncertainty for each group.

### Divergent patterns in phylogenetic clades and expected relationship disparity

Separate models for Mammalia-only, and Sauropsida-only were run (supplementary tables S33-40). Networks from sauropsidan indirect interactions did not differ from expected z-value (Est=- 0.28±0.93, 95% CI: -2.10 to 1.6) and sauropsidan direct networks did not differ from indirect networks (Est=-0.043±0.75, 95% CI: -1.50 to 1.40). Similarly, mammalian indirect networks did not differ from expected z-values (Est=0.10±1.09, 95% CI: -2.10 to 2.21) but direct interaction networks did significantly differ from mammalian indirect social interaction networks (Figure 4b; Est=2.04±0.72, 95% CI: 0.59 to 3.41). The posterior mean contribution of phylogeny was higher in the Sauropsida-only model (13.2%) than Mammalia-only model (3.1%) with specific species identity explaining 92.9% of variation in observed mammalian z-scores but only 41% in Sauropsida (supplementary tables S44, S46). When an overall phylogenetically controlled model was run with Mammalia and Sauropsida as a categorical variable, no significant difference was found between clade (Est=-0.23±0.95, 95% CI: -2.10 to 1.6) or between interaction types (Est=0.57±0.58, 95% CI: -0.58 to 1.7). In a further model where an interaction between categorical clade and social network interaction type was run, we found no significant interaction between clade and interaction type (Est=1.25±0.80, 95% CI: -0.33 to 2.83), hence this difference was only supported in Mammalia-only model but not in the Sauropsida-only model.

Together, these analyses show that smaller, sparser and more cohesive networks tend to exhibit stronger relationship disparity, that phylogeny and species identity drive a large part of the variation in observed relationship disparity and that this may reflect specific social structuring so that the disparity is weaker or stronger than expected from degree distribution and other structural properties alone.

## Discussion

We show here that the extent of the friendship paradox – the tendency for individuals to be connected to more highly connected partners than themselves (measured as relationship disparity) – is jointly shaped by network structure and biological context in wild vertebrate populations. Across 391 animal social networks, smaller and sparser networks with higher mean degree and greater group cohesion generally exhibited stronger relationship disparity. We find evidence of phylogenetic and specific species signals. Yet controlling for variation in individual sociability revealed that in mammalian networks saw a split with direct social interaction networks with a high disparity than would be expected from the network’s degree distribution, whereas sauropsidan networks did not dffer from expectations regardless of the interaction type of the network. These findings indicate that the strength of the Friendship Paradox in real animal populations reflects both the emergent consequences of the basic structural architecture of the network, and furthermore, may measure social or ecological processes that organise interactions between specific individuals.

Our results clarify how specific features of network structure contribute to local inequalities in connectedness. Relationship disparity increased with mean degree and decreased with edge density, indicating that it is not simply the total opportunity for interaction that matters, but how that opportunity is distributed. Social networks in which individuals have many potential ties but are far from fully connected, with high mean degree but relatively low density, provide more scope for some individuals’ neighbours to be substantially better connected than they are, thereby strengthening the friendship paradox. Once mean degree and edge density were accounted for, group cohesion was positively associated with relationship disparity, suggesting that cohesive subgroup structure amplifies differences between individuals and their neighbours in how deeply embedded they are in the wider network. Further, we also found that relationship disparity was only weakly related to degree heterogeneity and assortativity-by-centrality (see supplementary text), implying that it is not simply a re-labelling of familiar global metrics. Instead, it captures a higher-order, local property of social structure in how differences in degree are oriented between individuals and their immediate social associates.

Along with the network architecture, biological contexts were found to modify these structural effects in systematic ways. Direct and indirect social interaction networks did not differ strongly in their average relationship disparity after controlling for structural variables. For mammalian species only, they differed in how observed values compared to expectations based on variation in individuals’ sociability (i.e. permutation null models controlling for exact degree distributions), For mammalian networks based on indirect interactions (such as group membership or co-occurrence), observed relationship disparity was broadly consistent with that expected from their degree distributions and other structural properties, while social networks based on direct physical interactions showed systematically higher relationship disparity than expected, indicating greater uneven local social environments than their degree distributions alone would predict. These findings could reflect the trade-off that direct physical interactions have potentially greater costs than indirect non-physical interactions as direct physical contact poses a greater chance of interchange of microbes, such as pathogens or parasites, or risk of physical harm, whilst indirect interactions carry lower risks (27, 29–31). Direct interactions (e.g. mating, aggression), which may carry higher costs and benefits, might be more strongly shaped by social decisions (32), and may be constrained in ways that reproduce the degree-based expectation of relationship disparity rather than accentuating or suppressing it. On the other hand, networks based on indirect interactions, such as group membership, may be less limited and allow greater variation (8). As such approaches to disease mitigation such as contact tracing using the friendship paradox might work better than random sampling in indirect-based networks as the number of indirect connections is then expected to be linked to the rate of pathogen exposure (15, 33). As such, these findings add support to the hypothesis that social structure may itself be a product of social decisions and behaviour (1, 34, 35), and that the processes underlying the formation of social ties can lead to patterns of relationship disparity.

The phylogenetic differences in relationship disparity that we report here potentially point to broader variation in social network organisation across vertebrates. The evident phylogenetic signal and species variation may reflect the complex social systems across mammals vs non-mammalian vertebrates, often involving long-term bonds, dominance hierarchies, kin-biased associations, or differentiated social roles, and that these social levels can generate pronounced variation in social connectivity (32, 36). However, interestingly, relationship disparity values in sauropsidan social networks tended not to be greater than expected given their degree distribution. This suggests that the relationship disparity observed in sauropsidan systems may be largely driven by variation in individual sociability rather than other types of social structuring, which is consistent with relatively weak assortativity-by-degree in species such as great tits and social jays (37, 38). On the other hand, mammalian direct networks had significantly stronger relationship disparity than expected given their degree distributions compared to mammalian indirect networks. In mammals, this pattern could arise if these systems are likely to show strong assortativity by degree (hence reduced disparity as individuals connect based on sociability) or if behavioural or temporal constraints prevent a small subset of individuals from monopolising connections overall. In Sauropsida, both direct and indirect networks tended to have visually weaker relationship disparity strength than mammalian, which suggests relatively uniform local social environments which could potentially be related to lower cognitive capacity for social relationships (39–41) (but see (42–44)), or may partly reflect which reptile species and contexts are currently represented in social network repositories. While our findings provide evidence of taxonomic patterns in relationship disparity across systems, disentangling whether these primarily reflect cognitive differences, ecological constraints, or sampling biases will require further species-level analyses with richer behavioural data (45, 46).

Social relationships arise from patterns of interaction shaped by individual choices and constraints, reflecting trade-offs in forming and maintaining social connections. The structural and biological patterns identified here may have implications for how individuals experience their social environments, and for how processes that depend on social contact unfold. Relationship disparity captures a key aspect of individuals’ local social environment, by indicating whether individuals are surrounded by partners who are, on average, more, equally, or less socially connected than themselves. Relationship disparity may affect social decisions, for instance as individuals often adjust their number of connections based on observing the interactions of others, i.e. having an audience or third-party presence (47–49).

Relationship disparity at the individual level may also influence access to information and social support, exposure to pathogens, and opportunities for cooperation or conflict. An individual’s local social environment (defined by the social partners they are directly connected to) can have direct consequences for their fitness, by shaping survival and reproductive success, as demonstrated across diverse taxa including zebra finches (50), ravens (51), great tits (52, 53), red-legged partridge (54), primates (55, 56), meerkats (57), bottle-nosed dolphins (58), feral horses (59), cichlids (60), mice (61). The presence of observers can modify social behaviour even without direct involvement. In this context, relationship disparity can be interpreted as a measure of ‘social audience’ or third-party structure and may help contribute to our understanding of social processes, such as those in which individuals whose partners are highly connected occupy neighbourhoods where many others may observe, transmit information about, or mediate their interactions (47–49). By extending analysis beyond dyadic interactions, future work using relationship disparity may provide further insights into our understanding of individuals’ social experience and their resulting behavioural strategies in complex systems, over and above that coming from simply measuring their own connections (62–64).

As well as having implications for individual-level outcomes of relationship disparities, our findings may also hold relevance for applications that exploit the friendship paradox for surveillance and intervention (13, 14). Strategies that recruit the friends of randomly selected individuals, or that preferentially sample individuals whose neighbours are more connected than they are, have been used to detect infectious disease outbreaks earlier, to monitor behavioural change, and to target public health interventions more efficiently than random sampling (14, 15, 19, 20, 22–24). The added-value performance of these strategies depends essentially on both the extent of the friendship paradox, and whether it arises solely from degree heterogeneity, or from additional social structuring. Crucially for zoonotic surveillance, we show that more than 40% of animal networks have relationship disparity values that are significantly higher or lower than expected from their degree distributions, sizes and densities alone, and that deviations from null expectations depend on social interaction type, phylogeny and species identity. As such, in systems where relationship disparity is strong and close to null expectations (such as many sauropsidan networks and those based on indirect interactions) local information about ‘friends of friends’ may closely track global structure, making friendship-paradox-based sampling particularly powerful. By contrast, in systems where relationship disparity is stronger than expected (such as mammalian and direct social interaction networks) friendship-paradox strategies may not add value, or may require augmentation (for example, with spatial or temporal information, or with knowledge of social roles) to achieve similar gains in early detection or intervention efficiency.

Together, our comparative analysis provides a new empirical analysis of the Friendship Paradox in real-world animal systems and demonstrates how relationship disparity can be employed as a metric that complements other conceptual considerations of social structure, linking higher-order network architecture to biological context. The Friendship Paradox could also be extended beyond connectedness to individual attributes (for example, traits related to dominance) as in the generalised or attribute-based Friendship Paradox (4, 6, 65). Future developments could now enhance the implications of our study further. As animal social network repositories expand, future studies may be able to expand on these analyses by using high-resolution temporal networks across contexts (instead of time-aggregated ones) (66), separate networks into diverse interaction types (e.g. comparing across mating, aggression, grooming etc), assess more completely across taxa, and even integrate individual-level node information (sex, age etc). Such advances would be useful for examining whether individual-level differences in the friendship paradox predict changes in contagion risk, information flow or social stability. Answering these questions can shed light on how social systems evolve, how population declines might affect social structure, and lead to effective wildlife population management. As such, we call for broader consideration of relationship disparity within animal study systems, alongside other metrics, to help distil general principles governing the evolution and maintenance of animal societies.

## Materials and Methods

### Network selection

ASNR networks (28) were filtered to a set of criteria to fit the research questions. Networks classified as representing spatial proximity (n = 146) were removed, as we aimed to examine behavioural social associations and not only spatial positioning. As this research is based on natural/wild populations, networks that had a ‘captive’ population type were removed. Networks with fewer than 10 individuals and with more than 50% of nodes isolated with no connections were removed, as well as networks with an edge density of 1 as it is theoretically impossible to observe the friendship paradox on such networks.

We specified ‘Interaction type’ to represent the broad type of social interactions forming the network. Due to smaller sample sizes in some interaction type categories, interaction type was aggregated into direct physical contact and indirect non-physical contact interactions (see supplementary text). Networks of Amphibia, Arachnida, Insecta and Cephalopoda were removed due to small sample size of networks (of 2, 1, 1, 3 respectively), leaving 3 taxonomic classes: Aves, Mammalia and Non-avian Reptilia (n= 62, 283, 46, respectively), referred to as reptiles. The final dataset used consisted of 391 social networks of 53 vertebrate species. Due to 2 species of Reptilia this was combined with Aves as Sauropsida. All networks were treated as symmetrical (i.e. as undirected social associations).

### Relationship disparity calculations

We calculated ‘relationship disparity’ as a network-level measure to capture the local Friendship Paradox, by averaging local node-level relationship disparity across a network. In unweighted networks, connectedness was measured as degree. For each individual, relationship disparity was calculated as the mean degree of its direct neighbours minus the focal individual’s own degree. In weighted networks, connectedness was measured as strength, or weighted degree. For each individual, weighted relationship disparity was calculated as the mean strength of its direct neighbours minus the focal individual’s own strength. Thus, in weighted networks, edge weights are incorporated through each node’s total weighted degree, rather than by weighting each neighbour’s contribution to the neighbour mean. Positive values indicate that individuals are, on average, connected to social partners who are more connected than themselves; negative values indicate that individuals are, on average, more connected than their social partners. Therefore, the network-level relationship disparity is the mean of these individual-level values. The supplementary information provides the full equation, including how this local formulation is extended to weighted networks.

### Network calculations

The specific social network structure could enhance or decrease the extent of relationship disparity (i.e. the strength of the average level of Friendship Paradox). Three commonly used network metrics were calculated for each network: degree mean, edge density and group cohesion (supplementary figure S2) to capture different network structures. All measures were unweighted. Degree mean is the average number of connections, degree, for all nodes in the network and hence represents how many interactions an individual has with others. Edge density is a measure of the number of actual edges out of the theoretical maximum possible number of edges. It reflects how densely connected the network is, despite the network size and number of connections. Group cohesion is the minimum number of edges needed to remove to obtain a graph which is not strongly connected, and therefore functions as an intuitive modularity measure and shows how well integrated different sub-communities may be in a network. It is calculated from the ‘edge_connectivity’ function from ‘igraph’ package (67).

Two key biological features – interaction type and taxonomic class – likely to influence the structure of social networks and, consequently, relationship disparity were selected as biological predictors from within the ASNR database. Interaction type was aggregated in direct physical interaction and indirect associative networks (supplementary table S2).

### Phylogenetic Analysis

A phylogenetic tree depicting relationships among the constituent species in analysed social networks (figure 1) was created using Open Tree of Life data and package ‘rotl’ (68, 69) and plotted using packages ‘ggtree’ and ‘ggtreeExtra’ (70, 71). Two networks contained multiple species of the same taxonomic class, Aves. These were included in the phylogeny as the most common species in the original network (*Parus major* and *Acanthiza reguloides*). Grafen’s branch lengths computation (72) was used from the ‘ape’ package (73). The resulting ultrametric phylogenetic tree is on an arbitrary relative scale where all tips are equidistant from the root. To test for a phylogenetic signal in relationship disparity, a phylogenetic correlation matrix was included in each model where relevant, as a random effect. The correlation matrix was calculated using function ‘vcv’ from package ‘ape’ (73). Full results of models are shown in tables S6-19.

### Statistical analyses

We use Gaussian Bayesian phylogenetic mixed-effect models fit using the ‘brms’ package (74) to investigate how network and biological attributes predicted the strength of relationship disparity. All analysis were conducted in R version 4.3.1 (2023-06-16), using RStudio (75). Network analysis used ‘igraph’ (67) and the ‘tidyverse’ for data wrangling (76). As relationship disparity values across the networks were all positive and skewed, showing a power-law distribution, we used a natural log of relationship disparity as the response variable, modelled with a Gaussian distribution (see supplementary text)(77). To facilitate comparison of parameter estimates across models, continuous predictor variables were standardized (mean = 0, SD = 1). The size of the network and whether the network was weighted were controlled for all models as fixed effects. Weighted and unweighted networks were analysed in the same model as measures are similar except for the inclusion of weights (6).

Phylogenetic non-independence among species was modelled using a phylogenetic covariance matrix, species was included as a random effect to account for the inclusion of multiple networks per species in some cases. We first looked at network features and their association with relationship disparity. A model was fitted separately for degree mean, edge density and group cohesion, before fitting all these network features in the same model to see if they followed the same pattern when considered together in an additive model. Interaction type and taxonomic class were added separately as categorical variables. Finally, an overall model including all network features and interaction type was run to examine whether the same predictors were observed when all were considered simultaneously. Phylogenetic models were then run separately for only mammals (supplementary table S34-37) and only Sauropsids (supplementary table S38-41), with a respective sub-setted phylogenetic correlation matrix.

Results presented in figures were back-transformed for interpretability. Each model ran with four chains of 3,000 iterations each, with 1,000 iterations warmup and adapt delta of 0.95 to minimise divergent transitions. All models converged and had a R-hat of 1. Separate models with only species as a random effect, and models with species categorised into taxonomic class where also ran and detailed in the supplementary text. We used weakly informative priors for all model parameters. All fixed effects assigned normal priors (Normal(0,1)) and the random effects of phylogeny, species identity and residual error term has exponential(1) priors.

### Variance decomposition

Variance decomposition was performed to extract the components attributed to phylogeny, species identity, explanatory variables and residual variation. First, posterior draws were extracted from the models and variance was calculated for the for phylogeny, species identity, residual variation components (supplementary figure S4). The mean, lower and upper for the 95% confidence interval were then calculated and depicted in supplementary tables S41-46. Bayesian R2 was calculated using ‘bayes_r2’ from ‘rstantools’ (78) and is calculated according to Gelman et al. 2019 (79).

### Permutation testing

The specific degree distribution of a network may give rise to observed relationship disparity values. To examine whether these values were stronger or weaker than expected under the exact degree distribution of each network, permutation testing can be used. If the observed values are much greater or less than the expected, then the value, or network is described as ‘accentuated’ (see supplementary figure S3).

For each observed network, 1,000 network permutations were generated by rewiring the binary support graph while preserving the number of nodes and the exact binary degree sequence. Self-loops and multiple edges were prohibited. This preserves the number of edges, edge density, mean degree and number of isolated nodes. For weighted networks, the same binary degree-preserving rewiring was first applied, after which the observed set of edge weights was randomly reassigned across the permuted edges. This preserves the empirical distribution of edge weights, but not each node’s observed strength. This was intentional, because in weighted networks node strength is the connectedness quantity being compared. Preserving each node’s strength would therefore condition away a major component of the weighted social structure of interest.

Relationship disparity was then recalculated for each permuted network using the same calculation as was used for the corresponding observed network: degree-based relationship disparity for unweighted networks and strength-based relationship disparity for weighted networks. The observed value was compared to this network-specific null distribution, and a z-score was calculated as the difference between the observed value and the null mean, divided by the null standard deviation. These z-scores were interpreted as standardised, network-specific departures from null expectation: positive values indicate stronger relationship disparity than expected under the null model, whereas negative values indicate weaker relationship disparity than expected.

In addition, given that the z-score distribution is a standard normal distribution, we used Gaussian Bayesian mixed models to explore the z-scores calculated (see supplementary text). The permutation test controls the degree distribution, so only whether the network was weighted was included as a fixed effect. First, we looked at interaction type, then taxonomic class, then a model with both.

## Acknowledgments

EFN acknowledges funding from MRC (MR/W006731/1). JAF acknowledges funding from NERC (NE/V013483/2) and WildAI (C-2023-00057). SCLK acknowledges funding from the European Research Council under the European Union’s Horizon 2020 research and innovation programme (grant agreement no. 851550). We also thank R Mann for feedback, along with 2 anonymous reviews for useful comments.

