## Supplementary Information for "The Friendship Paradox across animal social systems is governed by network structure and biological features"

**Supporting Text**

**Relationship disparity calculations and expectations**

**1a) Description and Notations.** Here, we define the relationship disparity measures used in the manuscript and describe the structural expectations against which observed values were compared (1–5). Relationship disparity was calculated as a local, node-level, or ego-based form of the Friendship Paradox, and then averaged to obtain a network-level value. In unweighted networks, connectedness was measured as binary degree. In weighted networks, connectedness was measured as node strength, also referred to as weighted degree. Throughout the main text, both quantities are referred to as relationship disparity, with the corresponding unweighted and weighted-degree calculations specified below.

All networks were analysed as undirected social networks. Weighted networks were represented using non-negative edge weights. The weighted calculation used in the manuscript is a strength-based version of relationship disparity: it compares each node's own strength with the arithmetic mean strength of its direct neighbours. Thus, edge weights enter the calculation through node strength (not weighting each neighbour's contribution to the neighbour average).

**Table. S1.** Notation table for the explanation of relationship disparity calculations. Note that superscript $(s)$, as in $\delta_i^{(s)}$ and $\mathrm{RD}^{(s)}$, is a label for the strength-based version of the statistic (i.e. not an exponent). Similarly, subscripts such as $\mathrm{nz}$, $\mathrm{obs}$, and $\mathrm{null}$ are labels, not quantities being multiplied.

| Symbol | Meaning |
|---|---|
| $G = (V, E)$ | Undirected social network with node set $V$ and edge set $E$. |
| $n$ | Number of nodes in the network. |
| $m$ | Number of undirected edges in the binary support graph. |
| $A_{ij}$ | Binary adjacency matrix entry; $A_{ij} = 1$ if nodes $i$ and $j$ are connected, and $A_{ij} = 0$ otherwise. |
| $W_{ij}$ | Non-negative observed edge weight between nodes $i$ and $j$, with $W_{ij} = 0$ when $A_{ij} = 0$. In unweighted networks, $W_{ij} = A_{ij}$. |
| $N(i)$ | Set of neighbours directly connected to node $i$. |
| $k_i$ | Binary degree of node $i$, i.e. the number of direct neighbours of node $i$. |
| $s_i$ | Strength, or weighted degree, of node $i$, i.e. the sum of all edge weights incident to node $i$. |
| $\bar{k}_{N(i)}$ | Mean binary degree of the direct neighbours of node $i$, defined for $k_i > 0$. This replaces the ambiguous $\mu_{N(i)}^{k}$ notation. |
| $\bar{s}_{N(i)}$ | Mean strength, or weighted degree, of the direct neighbours of node $i$, defined for $k_i > 0$. This replaces the ambiguous $\mu_{N(i)}^{s}$ notation. |
| $V_{\mathrm{nz}}$ | Set of non-isolated nodes, i.e. nodes whose degree is greater than zero. |
| $n_{\mathrm{nz}}$ | Number of non-isolated nodes, i.e. the number of nodes in $V_{\mathrm{nz}}$. |
| $\delta_i$ | Local unweighted relationship disparity of node $i$. |

| Symbol | Meaning |
|---|---|
| $\delta_i^{(s)}$ | Local weighted-degree relationship disparity of node $i$, where the node attribute being compared is strength. |
| $\mathrm{RD}$ | Network-level unweighted relationship disparity, calculated as the mean of local unweighted values over all $n$ nodes after assigning isolates zero local disparity. |
| $\mathrm{RD}^{(s)}$ | Network-level weighted-degree relationship disparity, calculated as the mean of local weighted-degree values over all $n$ nodes after assigning isolates zero local disparity. |

**1bi) Unweighted relationship disparity**
For an unweighted network, connectedness is degree. The degree of node $i$ is

$$k_i = \sum_{j \in V} A_{ij}$$

For a node with $k_i > 0$, the mean degree of its direct neighbours is

$$\bar{k}_{N(i)} = \frac{1}{k_i} \sum_{j \in N(i)} k_j = \frac{1}{k_i} \sum_{j \in V} A_{ij}\, k_j$$

The first expression sums directly over the neighbour set $N(i)$. The second expression is the equivalent adjacency-matrix form: $A_{ij}$ acts as an indicator that includes node $j$ only when $j$ is a neighbour of $i$. The local relationship disparity of node $i$ is therefore

$$\delta_i = \bar{k}_{N(i)} - k_i = \frac{1}{k_i} \sum_{j \in V} A_{ij}\, k_j - k_i, \qquad k_i > 0$$

For isolated nodes, $k_i = 0$ and $\delta_i$ was set to 0 (see implementation note below). The network-level relationship disparity used in the manuscript is then

$$\mathrm{RD} = \frac{1}{n} \sum_{i \in V} \delta_i$$

A positive local value means that the focal individual is connected to neighbours who, on average, have more social partners than the focal individual. A negative local value means that the focal individual has more social partners than its neighbours on average. Thus, the node-level measure captures the local social environment experienced by each individual.

**1bii) Edge-based form of the unweighted measure**
For undirected unweighted networks, network-level relationship disparity can be rewritten as a sum over edges. Here, $\{i, j\} \in E$ denotes an unordered undirected edge, so each edge is counted once. Starting from the node-level definition,

$$n\,\mathrm{RD} = \sum_{i \in V_{\mathrm{nz}}} \left( \frac{1}{k_i} \sum_{j \in V} A_{ij}\, k_j - k_i \right)$$

Because each undirected edge $\{i, j\}$ contributes once from the perspective of $i$ and once from the perspective of $j$,

$$n\,\mathrm{RD} = \sum_{\{i,j\}\in E} \left(\frac{k_j}{k_i} + \frac{k_i}{k_j} - 2\right)$$

Equivalently,

$$n\,\mathrm{RD} = \sum_{\{i,j\}\in E} \frac{\left(k_i - k_j\right)^2}{k_i k_j}$$

and therefore

$$\mathrm{RD} = \frac{1}{n} \sum_{\{i,j\}\in E} \frac{\left(k_i - k_j\right)^2}{k_i k_j}$$

This edge-based expression clarifies what the unweighted relationship disparity measure captures. Under this definition, the network-level unweighted value cannot be negative in an undirected network, even though individual nodes may have positive or negative local values. This is because the network-level expression is a sum of edge-level degree differences. Edges connecting nodes with similar degree contribute little or nothing, whereas edges connecting nodes with very different degrees contribute more. Relationship disparity is therefore zero when connected nodes have equal degree throughout the network (including in complete graphs, regular graphs, and disconnected unions of degree-homogeneous components).

**1ci) Weighted-degree relationship disparity**

For weighted networks, connectedness was measured as node strength, also known as weighted degree. Node strength is the sum of the weights of all edges incident to the node:

$$s_i = \sum_{j\in V} W_{ij}$$

The weighted-degree version of relationship disparity compares the focal node's own strength with the mean strength of its direct neighbours. For a node with $k_i > 0$,

$$\bar{s}_{N(i)} = \frac{1}{k_i} \sum_{j\in N(i)} s_j = \frac{1}{k_i} \sum_{j\in V} A_{ij}\, s_j$$

The local weighted-degree relationship disparity is

$$\delta_i^{(s)} = \bar{s}_{N(i)} - s_i = \frac{1}{k_i} \sum_{j\in V} A_{ij}\, s_j - s_i, \qquad k_i > 0$$

For isolated nodes, $\delta_i^{(s)}$ was set to 0. The weighted-degree network-level relationship disparity is

$$\mathrm{RD}^{(s)} = \frac{1}{n} \sum_{i\in V} \delta_i^{(s)}$$

This is the calculation used for weighted networks in the manuscript. It asks whether an individual's direct social partners are, on average, more strongly connected overall than the individual itself. Edge weights enter the calculation through each node's strength (i.e weighted degree), $s_i$. Once strength has been calculated, however, the neighbour mean is an arithmetic mean over the focal node's neighbour set $N(i)$. In other words, each direct neighbour contributes once to the neighbour average, and the quantity being averaged is that neighbour's total weighted degree.

In the weighted-degree calculation used here, edge weights first contribute to each node's strength, $s_i$. Relationship disparity is then calculated by comparing each node's own strength with the arithmetic mean strength of its direct neighbours. Thus, each direct neighbour contributes once to the neighbour average, and the quantity being averaged is that neighbour's total strength. The calculation is therefore strength-based, rather than an edge-weighted neighbour average in which neighbours connected by stronger ties to the focal node would contribute more.

When all edge weights are equal to one, $W_{ij} = A_{ij}$ and $s_i = k_i$. In that special case, the weighted-degree expression reduces exactly to the unweighted relationship disparity expression above.

**1cii) Edge-based form of the weighted-degree measure**
The weighted-degree measure can also be written in an edge-based form. Again, $\{i,j\} \in E$ denotes an unordered undirected edge, so each edge is counted once. Starting from the node-level definition,

$$n\,\mathrm{RD}^{(s)} = \sum_{i \in V_{\mathrm{nz}}} \left( \frac{1}{k_i} \sum_{j \in V} A_{ij}\, s_j - s_i \right)$$

Rearranging the first term as a sum over undirected edges gives

$$n\,\mathrm{RD}^{(s)} = \sum_{\{i,j\} \in E} \left( \frac{s_j}{k_i} + \frac{s_i}{k_j} \right) - \sum_{i \in V} s_i$$

Because the sum of node strengths in an undirected weighted network is twice the total edge weight,

$$\sum_{i \in V} s_i = 2 \sum_{\{i,j\} \in E} W_{ij}$$

this becomes

$$\mathrm{RD}^{(s)} = \frac{1}{n} \sum_{\{i,j\} \in E} \left( \frac{s_j}{k_i} + \frac{s_i}{k_j} - 2W_{ij} \right)$$

This expression shows that the weighted-degree measure depends on three quantities: the strength of each node, the binary degree of the nodes across which neighbour averages are taken, and the weights of the edges joining those nodes.

With network-level weighted-degree relationship disparity, positive values indicate that, on average, individuals' neighbours have greater strength than the focal individuals themselves while negative values indicate the opposite. Because weighted-degree relationship disparity is measured on the scale of the edge weights, uniformly multiplying every weight in a network by a positive constant would multiply $\mathrm{RD}^{(s)}$ by the same constant. The permutation-based z-scores described below are not affected by this uniform rescaling, because both the observed value and the corresponding null distribution scale together. Raw weighted-degree relationship disparity values should therefore be interpreted on the scale of the weights used to construct each empirical network, whereas the null-model z-scores provide scale-standardised, network-specific departures from expectation.

*Implementation note:* Throughout our analyses, isolated nodes have no observed neighbours and therefore no defined neighbour average. As such, in the final implementation used for the analyses, isolated nodes were assigned local relationship disparity equal to zero before calculating the network mean. This treats isolated individuals as contributing no local neighbour disparity rather than leaving the network-level statistic undefined. Therefore, the network-level means below use denominator $n$, while only non-isolated nodes contribute non-zero terms. Networks with more than 50% isolated nodes were excluded from the main analyses. If one instead wanted the mean relationship disparity among non-isolated individuals only, the same formulae would use denominator $n_{\mathrm{nz}}$ rather than $n$. The statistic used in the manuscript is the all-node version with isolates contributing zero. Thus, for unweighted networks,

$$\mathrm{RD} = \left(\frac{n_{\mathrm{nz}}}{n}\right) \mathrm{RD}_{\mathrm{nz}}$$

where $\mathrm{RD}_{\mathrm{nz}}$ denotes the corresponding mean over non-isolated nodes only. The same scaling applies to the weighted-degree statistic:

$$\mathrm{RD}^{(s)} = \left(\frac{n_{\mathrm{nz}}}{n}\right) \mathrm{RD}_{\mathrm{nz}}^{(s)}$$

**1d) Structural interpretation and null expectations**

The calculations above (Sections 1a-1c) define relationship disparity as a local, node-level measure of how each individual's connectedness compares with that of its direct neighbours. Here, we first clarify how this local measure relates to established local and global formulations of the Friendship Paradox (Section 1di), and to related network structural descriptors such as degree heterogeneity, assortativity, mean degree, edge density, and network size (Section 1dii). We then describe the permutation-based null models used to compare observed relationship disparity with network-specific structural expectations, first for unweighted networks (Section 1diii), then for weighted networks (Section 1div), before explaining how null-model z-scores were calculated and interpreted (Section 1dv). Together, these sections distinguish the component of relationship disparity expected from basic network structure from additional organisation in how individuals are connected to one another.

**1di) Local and global formulations of the Friendship Paradox**

While the Friendship Paradox can be summarised using different, but related, formulations, we use a the local, or ego-based, formulation of relationship disparity. i.e. for each individual, we compare that individual's own connectedness with the mean connectedness of its direct neighbours and then average those local values across the network (giving each individual one contribution to the network-level estimate). In the global formulation, one compares the mean degree of a randomly selected node with the expected degree of a node reached by following a randomly selected edge. For unweighted networks, that global difference is determined by the degree distribution i.e. it is equal to the degree variance divided by the mean degree, provided that mean degree is greater than zero. Hence, degree heterogeneity is therefore central to the global Friendship Paradox as if all nodes have the same degree, the variance in degree is zero and the global Friendship Paradox disappears.

The local measure we use here also depends on how individuals with different connectedness values are arranged across edges. This distinction is closely related to recent theoretical work on inversity (4), which formalises how local Friendship Paradox patterns can vary among networks with the same degree distribution but different edgewise arrangements of degree. We do not analyse inversity as a separate empirical metric (or predictor, or response variable) in this study, but we cite this work to clarify why two networks with the same degree sequence can nevertheless differ in local relationship disparity, and we refer to inversity only as theoretical context for interpreting the local relationship disparity measure i.e. the degree sequence defines the connectedness values present in the network, whereas the arrangement of those values across edges determines whether individuals are connected to neighbours with similar or dissimilar levels of connectedness. For weighted networks, the same local interpretation applies, but the connectedness quantity being compared is node strength rather than binary degree. The weighted-degree calculation therefore asks whether an individual's direct social partners are, on average, more strongly connected overall than the individual itself. This is a local strength-based comparison (not a separate global attribute-based Friendship Paradox calculation).

**1dii) Relationship to degree heterogeneity, assortativity, and basic network structure**
As presented in the manuscript, and further in the Supplementary Information, relationship disparity is related to, but not identical to, degree heterogeneity or degree assortativity. Degree heterogeneity describes how much connectedness varies among individuals, without specifying which individuals are connected to which others, while degree assortativity describes whether individuals tend to connect to others with similar or dissimilar degree. Relationship disparity combines these ideas at the local level by asking whether individuals' direct neighbours are more or less connected than the focal individuals themselves.
Several basic network descriptors used in the manuscript are also mathematically related (but not identical). Referring to notation (see Section 1a - with $n$ nodes, $m$ edges, and edge density $d_E$), and considering a simple undirected graph, mean degree satisfies:

$$d_E = \frac{2m}{n(n-1)}, \qquad \bar{k} = \frac{2m}{n} = d_E(n-1).$$

As such, mean degree, edge density, and network size should not be interpreted as independent explanatory mechanisms, and in this research, these variables are treated as empirical structural descriptors and control variables that characterise where each observed animal social network lies within network-structural space by capturing basic differences in network size and connectivity among the empirical networks, and providing a basis for assessing whether observed relationship disparity is greater or lower than expected from these structural features. Network size, density, and mean degree provide useful context for interpreting relationship disparity because they describe basic constraints on the possible pattern of connections in a network. For example, as edge density approaches one, there is less scope for variation in degree among individuals; in a complete graph, all nodes have degree (n-1), and unweighted relationship disparity is zero. However, a higher mean degree does not necessarily imply stronger relationship disparity. A regular graph can have high mean degree while still having zero unweighted relationship disparity. Relationship disparity therefore depends on both variation in connectedness among individuals and how those connectedness values are arranged across edges.

**1diii) Degree-preserving null expectations for unweighted networks**
Relationship disparity may vary across empirical networks partly because networks differ in their basic structure, including their size, density, and degree distribution. To account for these structural differences, we compared each observed unweighted network with a set of permutation-based null networks generated from the same empirical network. As such, for each observed unweighted network, we generated 1,000 permuted networks by rewiring the binary support graph while preserving the number of nodes and the exact binary degree sequence. Self-loops and multiple edges were not permitted. This procedure preserves the number of edges, edge density, mean degree, degree variance, and the number of isolated nodes. It also preserves the degree-distribution component of the standard global Friendship Paradox. The unweighted null model therefore compares the observed local relationship disparity with values expected under the same degree sequence. In doing so, the degree values present in the network are held fixed, while the arrangement of edges among individuals is allowed to vary. This comparison tests whether the observed pattern of connections produces higher or lower local relationship disparity than expected from the degree sequence alone.

**1div) Weighted-network null expectations**
For weighted networks, we used the same binary degree-preserving rewiring procedure described above, and then randomly reassigned the observed edge weights across the permuted edges. This approach preserves the number of partners each individual has and the overall

distribution of edge weights in the network. The weighted null model was designed to evaluate weighted-degree relationship disparity relative to these structural features. In the observed networks, node strength depends on both the number of partners an individual has and the weights of those social ties. By preserving the binary degree sequence and the empirical edge-weight distribution, while allowing the allocation of weights to vary, the null model asks whether the observed pattern of weighted ties produces higher or lower weighted-degree relationship disparity than expected under these constraints.

**1dv) Null-model z-scores and interpretation**
Relationship disparity was recalculated for each permuted network using the same formula applied to the corresponding observed network: degree-based relationship disparity for unweighted networks and weighted-degree relationship disparity for weighted networks. For each empirical network with non-zero variance among its permuted values, we calculated a network-specific z-score as:

$$z = \frac{\mathrm{RD_{obs}} - \overline{\mathrm{RD}}_{\mathrm{null}}}{\mathrm{sd}(\mathrm{RD_{null}})}.$$

In this equation, RD denotes the relevant relationship-disparity statistic for that network: RD for unweighted networks and $\mathrm{RD}^{(s)}$ for weighted networks. Positive z-scores indicate that observed relationship disparity was greater than expected under that network's null model, whereas negative z-scores indicate that observed relationship disparity was lower than expected. The sign of the z-score therefore reflects the position of the observed value relative to the network-specific null distribution. These z-scores are best interpreted as standardised, network-specific departures from null expectation. As such, these z-scores provide standardised, network-specific comparisons between observed relationship disparity and the values expected under the permutation procedure. By using each empirical network as its own reference point, the null-model analysis assesses whether observed animal social networks show stronger or weaker relationship disparity than expected given their own structural constraints.

**1e) Note on scope of inference for comparative animal social networks**
The calculations and null models described in this section provide the mathematical basis for interpreting relationship disparity in the empirical animal social networks analysed in the manuscript. They are not intended to describe relationship disparity across all possible networks or graph classes. Instead, they define how relationship disparity was calculated for the comparative dataset used here, and how observed values were compared with network-specific structural expectations. Indeed, the empirical networks analysed in the manuscript represent real animal social networks from wild populations, drawn from comparative repositories and filtered using the inclusion criteria described in the main Methods. These networks differ in features such as species, interaction type, network size, density, weighting, and sampling method. This variation is relevant because relationship disparity can depend on both the structure of a network and how connectedness values are arranged across social ties. The calculations and null models described here therefore provide a framework for interpreting relationship disparity in the kinds of empirical animal social networks used to study processes such as social transmission, information flow, and group structure.
The analytical framework in this section therefore has three linked parts. First, observed relationship disparity is quantified using a local, node-level measure. Second, the measure is interpreted in relation to network structure, including degree, density, size, and edge weighting. Third, each empirical network is compared with a null expectation based on its own structural constraints. Together, these steps connect the Friendship Paradox basis of relationship disparity to the empirical variation observed across animal social networks.

References for calculations and structural explanations are found (1–5).

**Distribution choice for Bayesian models.** A gaussian distribution was chosen for the strength models due to the relationship disparity values being natural log transformed to be normal. A subsequent Shapiro-wilk test for normality (W = 0.09853, p-value = <2.2e -16), suggesting data deviates from normality. A student-t distribution was subsequently considered due to the large tails. On balance, we chose Gaussian for simplicity and interpretability because visual posterior predictive checks showed it captured the central body of data adequately; we verified sensitivity to Student-t and results were unchanged. A gaussian distribution was kept for permutated networks given the expected normal distribution of results.

**Interaction Type Category Aggregation.** Interaction type was aggregated into either direct physical or indirect non-physical interaction networks. Direct physical interactions were considered where contact between individuals must have occurred e.g. mating or grooming. Non-physical interactions were considered indirect interactions as no physical contact was presumed to have occurred, for example group membership. This interpretation is based on the ASNR classification of a network's interaction type. When category from ASNR was not clear, individual papers were read to correctly classify e.g. dominance. Full definitions of original interaction types are described in Sah *et al.* 2019 and in Table S4 (6).

**Table. S2.** Interaction type category aggregation with sample size

| Interaction type (aggregated) | No. Networks | Interaction type (original) | No. Networks |
|---|---|---|---|
| Direct | 89 | FE_mating | 11 |
| | | dominance | 22 |
| | | food_sharing | 5 |
| | | grooming | 32 |
| | | group_membership | 6 |
| | | overall_mix | 3 |
| | | physical_contact | 8 |
| | | social_projection_bipartite | 2 |
| Indirect | 302 | group_membership | 49 |
| | | non_physical_social_interaction | 6 |
| | | social_projection_bipartite | 247 |

**Phylopic attribution for animal illustrations in main text figure 1.** Andrew A. Farke (*Gopherus agassizii*), Andy Wilson (*Elephas maximus*, *Molothrus ater*), Arcadia Science (*Mus musculus*), Archibald Thorburn (*Myotis mystacinus*), Chris huh (*Orcinus orca*), Edwin Price (*Gallinago stricklandii*), Gabriela Palomo-Munoz (*Ateles geoffroyi*, *Ovis canadensis*), Jonathan Lawley (*Pan troglodytes*), Kai Caspar (*Equus grevyi*, *Papio papio*), Kai R. Caspar (*Cercopithecus diana*, *Macaca sylvanus*), Lukasiniho (*Bos bison*), Margot Michaud (*Cryptoprocta ferox*, *Zalophus wollebaeki*), Matt Wilkins (*Hirundo rustica*), Matt Wilkins (photo by Patrick Kavanagh) (*Acanthiza lineata*), Michael Scroggie (*Macropus giganteus*), Oscar Sanisidro (*Crocuta crocuta*), Rachel T Mason (*Trichosurus vulpecula*), Steven Traver (*Calloselasma rhodostoma*, *Erythrocebus patas*), and others (*Alouatta caraya*, *Dipodomys*, *Haemorhous mexicanus mexicanus*, *Macaca mulatta*, *Saimiri*, *Tursiops truncatus*).

**Difference between relationship disparity, degree heterogeneity and degree assortativity**
Relationship disparity is different from both degree heterogeneity and degree assortativity. Degree heterogeneity is relative, whilst the relationship disparity is concerned with the absolute relative number of individuals and is not proportional to the size of the network. Degree assortativity is weakly associated with strength of relationship disparity, and with different extents depending on the subset of the data into weighted and unweighted datasets. Relationship disparity is an amalgamation of degree heterogeneity, degree assortativity and other structural features such as triadic closure and modularity. Because it is absolute, it is powerful local measure. Please refer to figure S1 and table S3.

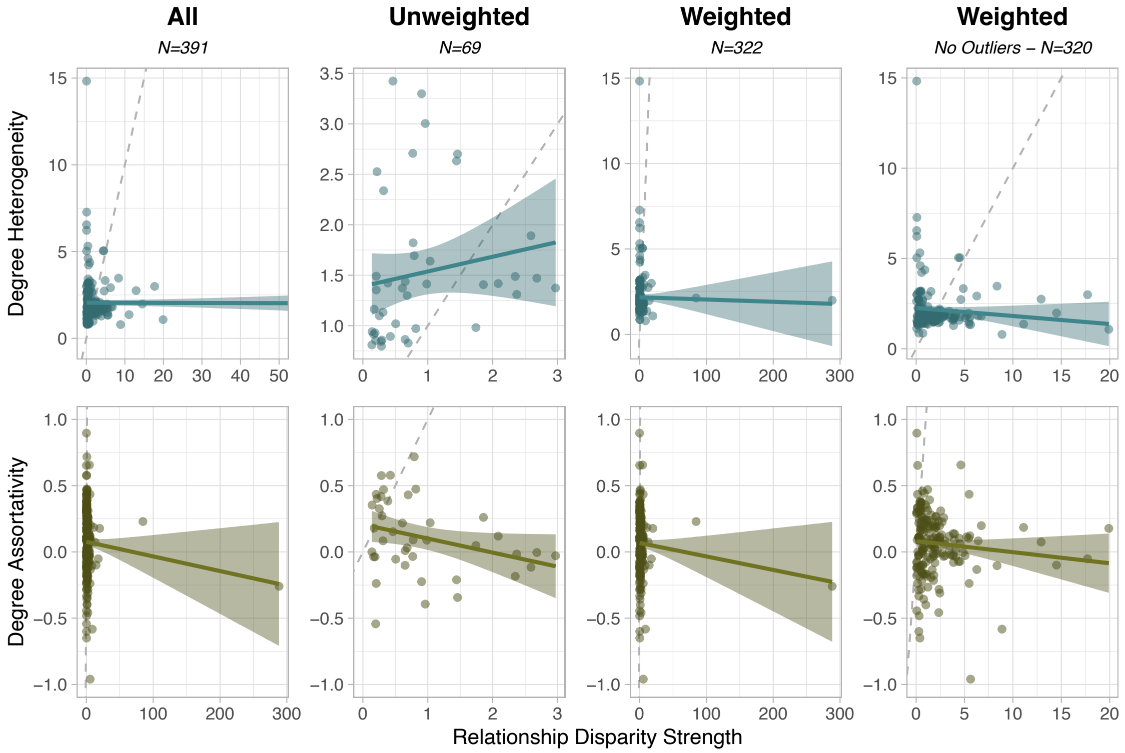


**Fig. S1.** Scatter plot of the relationship between strength of relationship disparity and degree heterogeneity (top) and degree assortativity (bottom), for all networks, unweighted, weighted, and weighted removing two outliers. Dashed line represents a line of y = x, and thus a 1:1 relationship between Relationship disparity and the network metric.

**Table. S3.** Linear regression models between relationship disparity and degree heterogeneity and degree assortativity.

| Linear model | Slope Estimate | Standard Error | T value | p Value | $r^2$ | Adjusted $r^2$ |
|---|---|---|---|---|---|---|
| Heterogeneity_all | -0.0005 | 0.0042 | -0.1199 | 0.9047 | 0.0001 | -0.0043 |
| Heterogeneity_unweighted | 0.1462 | 0.1402 | 1.0427 | 0.3033 | 0.0265 | 0.0021 |
| Heterogeneity_weighted | -0.0013 | 0.0044 | -0.2997 | 0.7647 | 0.0005 | -0.0049 |
| Heterogeneity_weighted_outliers | -0.0445 | 0.0345 | -1.2929 | 0.1976 | 0.0090 | 0.0036 |
| Assortativity_all | -0.0011 | 0.0008 | -1.3316 | 0.1843 | 0.0077 | 0.0034 |
| Assortativity_unweighted | -0.1062 | 0.0534 | -1.9897 | 0.0535 | 0.0901 | 0.0673 |
| Assortativity_weighted | -0.0084 | 0.0063 | -1.3362 | 0.1831 | 0.0096 | 0.0042 |
| Assortativity_weighted_outliers | -0.0084 | 0.0063 | -1.3362 | 0.1831 | 0.0096 | 0.0042 |

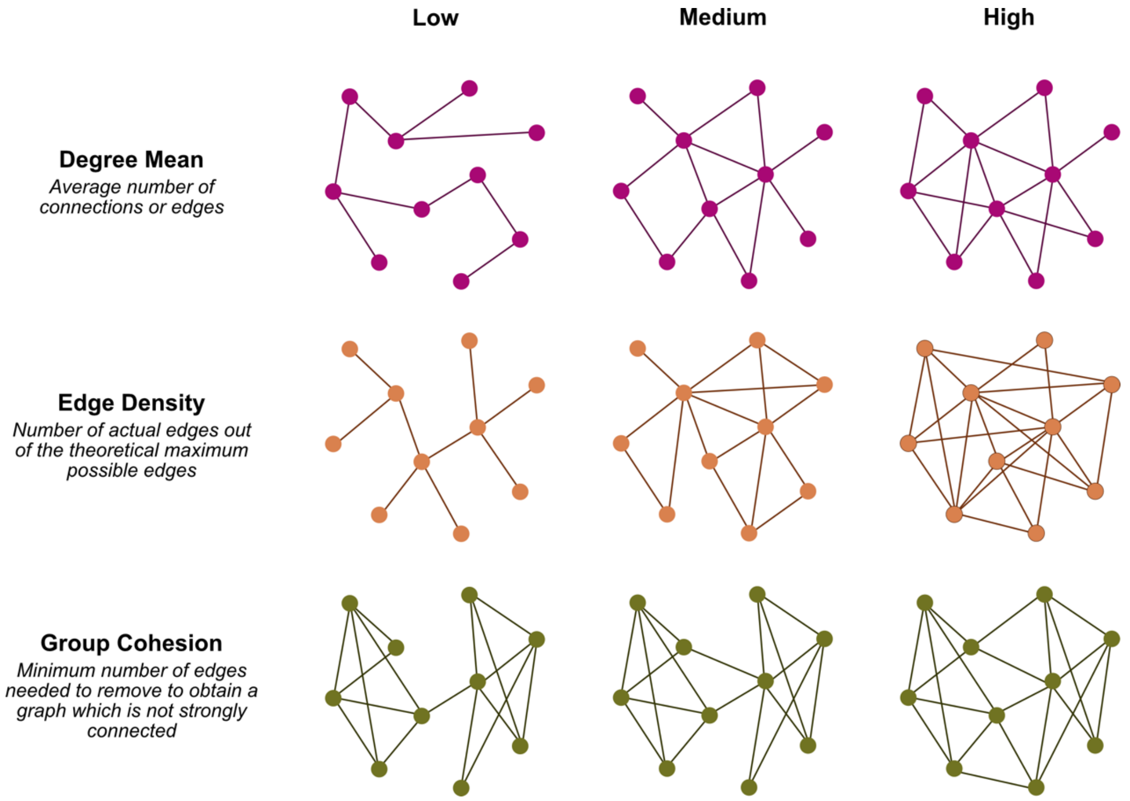


**Fig. S2.** Illustration of network metrics varying in strength**.** Degree mean (top), Edge density (middle) and Group cohesion/graph adhesion (bottom) for various strengths of network metric for a randomly generated network of 10 nodes.

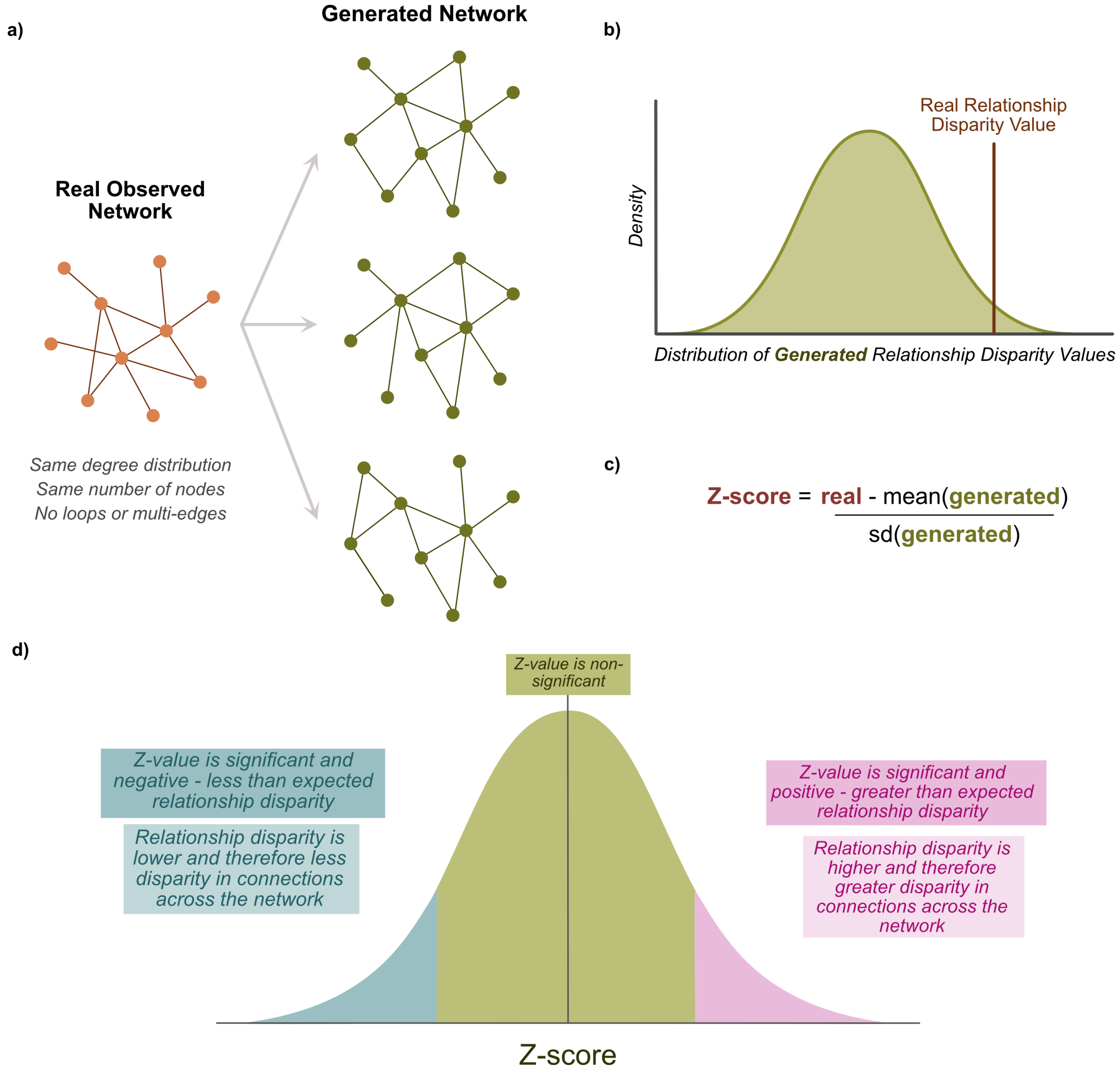


**Fig. S3.** Permutation test involved taking the same degree distribution and size of network then rewiring the network to generate networks with the same basic network features (a) whilst removing loops and multi-edges. Each of these networks' mean relationship disparity was calculated, and a distribution was calculated from these series of values (b). The real observed relationship disparity z-score was then calculated from this distribution of values mean and standard deviation (c). d) Values falling greater and less than the expected mean relationship disparity suggesting that in different social networks the relationship disparity is reduced or enhanced.

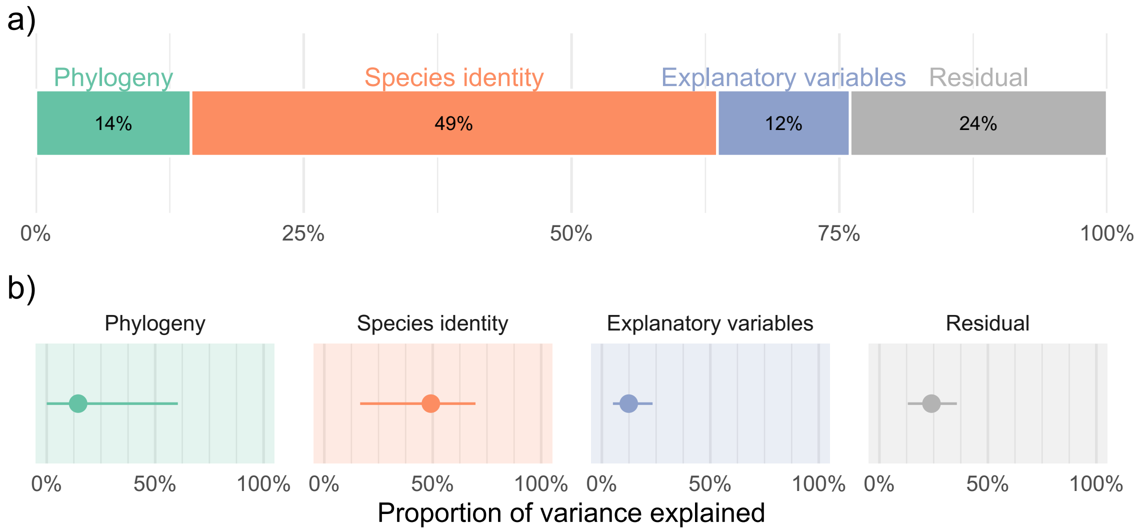


**Fig. S4** Variance decomposition of phylogenetically-controlled gaussian Bayesian model with formula of *meanlog ~ nodes_scale + weighted + degree_mean_scale + edge_density_scale + group_cohesion_scale + interaction_type + clade + (1 | gr(phylo, cov = cov_matrix)) + (1 | species_name).* a) is the mean variance across the different draws of the model. B) is the proportion of variance explained, where the points represent mean proportion of variance explained across all model draws with 95% credible interval for overall proportion values across all draws. Variance is attributed to shared phylogenetic history (green), species-specific identity (orange), explanatory variables in this case network features interaction type in blue and residual unexplained variance in grey. Full model output is in supplementary table S13.

**Table S4:** Dataset table describing species, taxonomic class, interaction type aggregated, original interaction type as described by ASNR, definitions of original interaction, number of networks for each and the citation for the network, based on ASNR.

| Species | Taxonomic Class | Interaction type | | | No. Networks | Citation |
|---|---|---|---|---|---|---|
| | | ***Aggregated*** | ***Original*** | ***Definition*** | | |
| Agkistrodon contortrix | Reptilia | Direct | FE_mating | fluid exchange occurs | 1 | Levine, B. A., C. F. Smith, G. W. Schuett, M. R. Douglas, M. A. Davis, and M. E. Douglas. 2015. Bateman_Trivers in the 21st Century: sexual selection in a North American pitviper. Biol. J. Linn. Soc. 114:436_445. |
| Alouatta caraya | Mammalia | Direct | FE_mating | fluid exchange occurs | 1 | Oklander, L. I., M. Kowalewski, and D. Corach. 2014. Male reproductive strategies in black and gold howler monkeys (Alouatta caraya). Am. J. Primatol. 76:43_58 |
| Argya squamiceps | Aves | Direct | dominance | brief aggressive and ritualised chases, usually towards a lower ranked individual. | 5 | Dragic, N., Keynan, O., & Ilany, A. (2021). Multilayer social networks reveal the social complexity of a cooperatively breeding bird. iScience, 24(11), 103336. https://doi.org/10.1016/j.isci.2021.103336 |
| Argya squamiceps | Aves | Direct | food_sharing | Allofeeding- act of an individual feeding another individual within the group. Can be an act of parental care or a dominance display | 5 | Dragic, N., Keynan, O., & Ilany, A. (2021). Multilayer social networks reveal the social complexity of a cooperatively breeding bird. iScience, 24(11), 103336. https://doi.org/10.1016/j.isci.2021.103336 |
| Argya squamiceps | Aves | Direct | grooming | Allopreening- an individual cleaning and grooming other members of the group | 5 | Dragic, N., Keynan, O., & Ilany, A. (2021). Multilayer social networks reveal the social complexity of a cooperatively breeding bird. iScience, 24(11), 103336. https://doi.org/10.1016/j.isci.2021.103336 |
| Argya squamiceps | Aves | Direct | physical_contact | play-fights, tug-of-war and displacements. Several play signals like crouching, rolling over, elevation of sticks, play bow, eye contact, freezing | 4 | Dragic, N., Keynan, O., & Ilany, A. (2021). Multilayer social networks reveal the social complexity of a cooperatively breeding bird. iScience, 24(11), 103336. https://doi.org/10.1016/j.isci.2021.103336 |
| Argya squamiceps | Aves | Indirect | non_physical_social_interaction | scrounging- direct theft, joining food patches discovered by others, taking over food patches discovered by others | 6 | Dragic, N., Keynan, O., & Ilany, A. (2021). Multilayer social networks reveal the social complexity of a cooperatively breeding bird. iScience, 24(11), 103336. https://doi.org/10.1016/j.isci.2021.103336 |
| Ateles geoffroyi | Mammalia | Direct | grooming | Based on observation of grooming interaction | 2 | Griffin, Randi H., and Charles L. Nunn. "Community structure and the spread of infectious disease in primate social networks." Evolutionary Ecology 26.4 (2012): 779-800. |

| | | | | | | |
|---|---|---|---|---|---|---|
| Ateles hybridus | Mammalia | Direct | physical_contact | Physical contact event between the two individuals (e.g. grooming, mating and embracing) | 1 | Rimbach, Rebecca, et al. "Brown spider monkeys (Ateles hybridus): a model for differentiating the role of social networks and physical contact on parasite transmission dynamics." Phil. Trans. R. Soc. B 370.1669 (2015): 20140110. |
| Bison bison | Mammalia | Direct | dominance | Only those aggressive interactions were analyzed that led to an outcome in which the lost and the winner could be categorized unambiguosly | 1 | Dale F Lott. Dominance relations and breeding rate in mature male American bison. Zeitschrift Tierpsychologie, 49(4):418-432, 1979. |
| Bos taurus | Mammalia | Direct | dominance | The dominance relationship was ascertained from direct physical contests. | 1 | Martin W. Schein and Milton H. Fohrman. Social dominance relationships in a herd of dairy cattle. The British J. of Animal Behaviour, 3(2):45-55, 1955. |
| Brachyteles arachnoides | Mammalia | Indirect | group_membership | Close social association less than 5m | 1 | Griffin, Randi H., and Charles L. Nunn. "Community structure and the spread of infectious disease in primate social networks." Evolutionary Ecology 26.4 (2012): 779-800. |
| Canis familiaris | Mammalia | Direct | FE_mating | fluid exchange occurs | 1 | Cafazzo, S., R. Bonanni, P. Valsecchi, and E. Natoli. 2014. Social Variables Affecting Mate Preferences, Copulation and Reproductive Outcome in a Pack of Free-Ranging Dogs. PLoS ONE 9 |
| Canis familiaris | Mammalia | Direct | FE_mating | fluid exchange occurs | 2 | Pal, S. K. 2011. Mating System of Free-Ranging Dogs (Canis familiaris). Int. J. Zool. e314216. |
| Cercopithecus campbelli | Mammalia | Direct | grooming | Based on observation of grooming interaction | 1 | Griffin, Randi H., and Charles L. Nunn. "Community structure and the spread of infectious disease in primate social networks." Evolutionary Ecology 26.4 (2012): 779-800. |
| Crocuta crocuta | Mammalia | Indirect | group_membership | Association patterns were recorded based on the co-occurrence of each pair of individuals, during the period for which they were concurrently present in the clan. | 3 | Holekamp, Kay E., et al. "Society, demography and genetic structure in the spotted hyena." Molecular Ecology 21.3 (2012): 613-632. |
| Cryptoprocta ferox | Mammalia | Direct | FE_mating | fluid exchange occurs | 1 | Hawkins, C. E., and P. A. Racey. 2009. A novel mating system in a solitary carnivore: the fossa. J. Zool. 277:196_204 |
| Dipodomys spectabilis | Mammalia | Direct | FE_mating | fluid exchange occurs | 1 | Randall, J. A. 1991. Mating strategies of a nocturnal, desert rodent (Dipodomys spectabilis). Behav. Ecol. Sociobiol. 28:215_220 |
| Elephas maximus | Mammalia | Direct | dominance | Indicators of dominance as well as subordination was included. If a series of interactions occurred during a particular event, the winners/losers were determined only on conclusion of the event, when individuals or groups moved apart. | 1 | de Silva, Shermin, Volker Schmid, and George Wittemyer. "Fission-fusion processes weaken dominance networks of female Asian elephants in a productive habitat." Behavioral Ecology (2016): arw153. |

| | | | | | | |
|---|---|---|---|---|---|---|
| Equus grevyi | Mammalia | Indirect | group_membership | A group was defined as a set of one or more individuals that is spatially cohesive and distinct from other groups at the time of observation. Edges were constructed based on half-weight index (HWI). | 1 | Siva R Sundaresan, Ilya R Fischhoff, Jonathan Dushoff, and Daniel I Rubenstein. Network metrics reveal differences in social organization between two fission-fusion species, Grevy's zebra and onager. Oecologia, 151(1):140-149, 2007. |
| Erythrocebus patas | Mammalia | Direct | grooming | Based on observation of grooming interaction | 2 | Griffin, Randi H., and Charles L. Nunn. "Community structure and the spread of infectious disease in primate social networks." Evolutionary Ecology 26.4 (2012): 779-800. |
| Gallinago media | Aves | Direct | FE_mating | fluid exchange occurs | 2 | Fiske, P., and J. A. K_l_s. 1995. Mate sampling and copulation behaviour of great snipe females. Anim. Behav. 49:209_219. |
| Gopherus agassizii | Reptilia | Indirect | social_projection_bipartite | A bipartite network was first constructed based on burrow use - an edge connecting a tortoise node to a burrow node indicated burrow use by the individual. Social networks of desert tortoises were then constructed by the bipartite network into a single-mode projection of tortoise nodes. | 45 | Sah, Pratha, et al. "Inferring social structure and its drivers from refuge use in the desert tortoise, a relatively solitary species." Behavioral Ecology and Sociobiology 70.8 (2016): 1277-1289. |
| Haemorhous mexicanus | Aves | Indirect | social_projection_bipartite | A machine learning algorithm was applied to identify clusters of detections on feeders. Next, the network was generated based on patterns of co-occurrence by individuals in the same feeding events. Asociations between birds were defined using the simple ratio index. | 1 | Adelman, James S., et al. "Feeder use predicts both acquisition and transmission of a contagious pathogen in a North American songbird." Proc. R. Soc. B. Vol. 282. No. 1815. The Royal Society, 2015. |
| Helogale parvula | Mammalia | Direct | grooming | all observed grooming bouts lasting longer than 5 s that occurred between individuals aged 12 months and older | 2 | Kern, J.M., Radford, A.N. Strongly bonded individuals prefer to forage together in cooperatively breeding dwarf mongoose groups. Behav Ecol Sociobiol 75, 85 (2021). https://doi.org/10.1007/s00265-021-03025-0 |
| Hirundo rustica | Aves | Direct | physical_contact | Interaction 0.1 m and closer | 2 | Levin, Iris I., et al. "Stress response, gut microbial diversity and sexual signals correlate with social interactions." Biology Letters 12.6 (2016): 20160352. |
| Macaca arctoides | Mammalia | Direct | grooming | Based on observation of grooming interaction | 2 | Griffin, Randi H., and Charles L. Nunn. "Community structure and the spread of infectious disease in primate social networks." Evolutionary Ecology 26.4 (2012): 779-800. |
| Macaca assamensis | Mammalia | Direct | grooming | Based on observation of grooming interaction | 1 | Griffin, Randi H., and Charles L. Nunn. "Community structure and the spread of infectious disease in primate social networks." Evolutionary Ecology 26.4 (2012): 779-800. |

| | | | | | | |
|---|---|---|---|---|---|---|
| Macaca assamensis | Mammalia | Direct | overall_mix | recorded all social interactions, including grooming, body contact, approaches into and departures from a distance of 1.5 m to other individuals and vice versa, and agonistic interactions | 1 | Puga-Gonzalez, I., Ostner, J., Schülke, O., Sosa, S., Thierry, B., & Sueur, C. (2018). Mechanisms of reciprocity and diversity in social networks: a modeling and comparative approach. Behavioral Ecology, 29(3), 745-760. |
| Macaca fuscata | Mammalia | Direct | grooming | Based on observation of grooming interaction | 1 | Griffin, Randi H., and Charles L. Nunn. "Community structure and the spread of infectious disease in primate social networks." Evolutionary Ecology 26.4 (2012): 779-800. |
| Macaca fuscata | Mammalia | Direct | grooming | grooming defined as a monkey picks through and/or slowly brushes aside the fur of another individual with one or both hands | 1 | Puga-Gonzalez, I., Ostner, J., Schülke, O., Sosa, S., Thierry, B., & Sueur, C. (2018). Mechanisms of reciprocity and diversity in social networks: a modeling and comparative approach. Behavioral Ecology, 29(3), 745-760. |
| Macaca fuscata | Mammalia | Direct | group_membership | intergroup encounter occurred when two groups approached and members of the study group were regarded to have recognized the other group | 1 | Puga-Gonzalez, I., Ostner, J., Schülke, O., Sosa, S., Thierry, B., & Sueur, C. (2018). Mechanisms of reciprocity and diversity in social networks: a modeling and comparative approach. Behavioral Ecology, 29(3), 745-760. |
| Macaca fuscata | Mammalia | Direct | dominance | The dominance relations between females were determined based on approach-retreat episodes around the food. The dominance range order was arranged based on these dyadic relations. | 1 | Takahata, Yukio. "Diachronic changes in the dominance relations of adult female Japanese monkeys of the Arashiyama B group." The monkeys of Arashiyama. State University of New York Press, Albany (1991): 123-139. |
| Macaca fuscata | Mammalia | Direct | dominance | based on approach retreat episodes around food and dyadic relations | 1 | Takahata, Y. (1991). Diachronic changes in the dominance relations of adult female Japanese monkeys of the Arashiyama B group. The monkeys of Arashiyama. State University of New York Press, Albany, 123-139. |
| Macaca fuscata | Mammalia | Direct | group_membership | intergroup encounter occurred when two groups approached and members of the study group were regarded to have recognized the other group | 1 | Puga-Gonzalez, I., Ostner, J., Schülke, O., Sosa, S., Thierry, B., & Sueur, C. (2018). Mechanisms of reciprocity and diversity in social networks: a modeling and comparative approach. Behavioral Ecology, 29(3), 745-760. |
| Macaca mulatta | Mammalia | Direct | grooming | Based on observation of grooming interaction | 1 | Griffin, Randi H., and Charles L. Nunn. "Community structure and the spread of infectious disease in primate social networks." Evolutionary Ecology 26.4 (2012): 779-800. |
| Macaca mulatta | Mammalia | Direct | grooming | observation of allo-grooming | 1 | Puga-Gonzalez, I., Ostner, J., Schülke, O., Sosa, S., Thierry, B., & Sueur, C. (2018). Mechanisms of reciprocity and diversity in social networks: a modeling and comparative approach. Behavioral Ecology, 29(3), 745-760. |
| Macaca radiata | Mammalia | Direct | grooming | Based on observation of grooming interaction | 2 | Griffin, Randi H., and Charles L. Nunn. "Community structure and the spread of infectious disease in |

| | | | | | | |
|---|---|---|---|---|---|---|
| | | | | | | primate social networks." Evolutionary Ecology 26.4 (2012): 779-800. |
| Macaca sylvanus | Mammalia | Direct | dominance | Aggressive behaviour ie. Bite, charge, chase, displace, grab, lunge, slap | 1 | Fedurek, Piotr, Richard McFarland, Bonaventura Majolo, and Julia Lehmann. 2022. 'Social Responses to the Natural Loss of Individuals in Barbary Macaques'. *Mammalian Biology* 102 (4): 1249–66. https://doi.org/10.1007/s42991-022-00283-x. |
| Macaca sylvanus | Mammalia | Direct | grooming | grooming behaviour (physical contact) | 1 | Fedurek, Piotr, Richard McFarland, Bonaventura Majolo, and Julia Lehmann. 2022. 'Social Responses to the Natural Loss of Individuals in Barbary Macaques'. *Mammalian Biology* 102 (4): 1249–66. https://doi.org/10.1007/s42991-022-00283-x. |
| Macaca sylvanus | Mammalia | Direct | group_membership | observed allogrooming and agonistic behaviors (threatening face or growl, charge, avoidance, attack, chase, and aggressive slap, grab, or bite) | 1 | Puga-Gonzalez, I., Ostner, J., Schülke, O., Sosa, S., Thierry, B., & Sueur, C. (2018). Mechanisms of reciprocity and diversity in social networks: a modeling and comparative approach. Behavioral Ecology, 29(3), 745-760. |
| Macaca tonkeana | Mammalia | Direct | group_membership | Contact sitting | 1 | Griffin, Randi H., and Charles L. Nunn. "Community structure and the spread of infectious disease in primate social networks." Evolutionary Ecology 26.4 (2012): 779-800. |
| Macropus giganteus | Mammalia | Direct | group_membership | Two individuals were assumed to be associating if they occured within 120 cm of another at set 15-min intervals in the enclosure. | 1 | TR Grant. Dominance and association among members of a captive and a free-ranging group of grey kangaroos (Macropus giganteus). Animal Behaviour, 21(3):449-456, 1973. |
| Microtus agrestis | Mammalia | Indirect | social_projection_bipartite | An edge was inserted into the network whenever two voles were caught in at least one common trap over the primary trapping sessions being considered | 174 | Davis, Stephen, et al. "Spatial analyses of wildlife contact networks." Journal of the Royal Society Interface 12.102 (2015): 20141004. |
| Mirounga angustirostris | Mammalia | Direct | dominance | Dominance status determined by Elo rating based on winner of an competitive interaction and intensity of interaction | 7 | Casey, Caroline, et al. "Rival assessment among northern elephant seals: evidence of associative learning during male-male contests." Royal Society open science 2.8 (2015): 150228. |
| mixed species (Acanthiza reguloides) | Aves | Indirect | group_membership | Flock membership was identified based on frequent interactions between (such as _beating¬ê for insects) flocks and large gaps between flocks. Association strength of each dyad was calculated using the simple ratio index. | 1 | Farine, Damien R., and Peter J. Milburn. "Social organisation of thornbill-dominated mixed-species flocks using social network analysis." Behavioral Ecology and Sociobiology 67.2 (2013): 321-330. |
| mixed species (Parus major) | Aves | Indirect | social_projection_bipartite | Groups were defined as individuals detected on the same nest-box during the same day, and co-memberships represented individuals | 6 | Firth, Josh A., and Ben C. Sheldon. "Experimental manipulation of avian social structure reveals segregation is carried over across contexts." |

| | | | | | | |
|---|---|---|---|---|---|---|
| | | | | that overlapped in nest-box exploration patterns during the same day. Networks were calculated from these group-by-individual matrices using the halfweight index. | | Proceedings of the Royal Society of London B: Biological Sciences 282.1802 (2015): 20142350. |
| Molothrus ater | Aves | Direct | FE_mating | fluid exchange occurs | 1 | Yokel, D. A. 1986. Monogamy and brood parasitism: an unlikely pair. Anim. Behav. 34:1348_1358. |
| Mus musculus domesticus | Mammalia | Indirect | social_projection_bipartite | information on identity of animal entering or exiting, nest box number, and timestamp were collected | 1 | Lopes, P. C., Block, P., & K_nig, B. (2016). Infection-induced behavioural changes reduce connectivity and the potential for disease spread in wild mice contact networks. Scientific reports, 6, 31790. |
| Myotis sodalis | Mammalia | Indirect | social_projection_bipartite | Roost network was first contructed as a two-mode network that consisted of bats and roosts. Single-mode projection of the bat nodes was used to assess colony social structure. | 1 | Silvis, Alexander, et al. "Roosting and foraging social structure of the endangered Indiana bat (Myotis sodalis)." PloS one 9.5 (2014): e96937. |
| Orcinus orca | Mammalia | Indirect | group_membership | recorded social interactions between individuals | 1 | Guimarães Jr, P. R., de Menezes, M. A., Baird, R. W., Lusseau, D., Guimaraes, P., & Dos Reis, S. F. (2007). Vulnerability of a killer whale social network to disease outbreaks. Physical Review E, 76(4), 042901. |
| Otospermophilus beecheyi | Mammalia | Direct | social_projection_bipartite | belowground associations determined using PIT tags and aboveground associations determined by observation | 2 | Smith, J. E., Gamboa, D. A., Spencer, J. M., Travenick, S. J., Ortiz, C. A., Hunter, R. D., & Sih, A. (2018). Split between two worlds: automated sensing reveals links between above-and belowground social networks in a free-living mammal. Philosophical Transactions of the Royal Society B: Biological Sciences, 373(1753), 20170249. |
| Ovis canadensis | Mammalia | Direct | dominance | Social status was determined by assembling a win-loss matrix based on the outcome of agonistic interactions. The winner of a dominance fight, involving a series of butts and clashes, was recorded as winning one interaction. | 2 | Christine C Hass. Social status in female bighorn sheep (Ovis canadensis): Expression, development and reproductive correlates. J. of Zoology, 225(3):509-523, 1991. |
| Pan paniscus | Mammalia | Direct | grooming | Based on observation of grooming interaction | 1 | Griffin, Randi H., and Charles L. Nunn. "Community structure and the spread of infectious disease in primate social networks." Evolutionary Ecology 26.4 (2012): 779-800. |
| Pan troglodytes | Mammalia | Direct | grooming | grooming behaviour (physical contact) | 1 | Rawlings, Bruce S., Edwin J. C. Van Leeuwen, and Marina Davila-Ross. 2023. 'Chimpanzee Communities Differ in Their Inter- and Intrasexual Social Relationships'. *Learning & Behavior* 51 (1): |

| | | | | | | |
|---|---|---|---|---|---|---|
| | | | | | | 48–58. https://doi.org/10.3758/s13420-023-00570-8. |
| Pan troglodytes | Mammalia | Direct | grooming | Based on observation of grooming interaction | 1 | Griffin, Randi H., and Charles L. Nunn. "Community structure and the spread of infectious disease in primate social networks." Evolutionary Ecology 26.4 (2012): 779-800. |
| Papio anubis | Mammalia | Direct | grooming | grooming behaviour (physical contact) | 2 | Pritchard, Alexander J., et al. ‚Individual Differences in Coping Styles and Associations with Social Structure in Wild Baboons (Papio Anubis). Animal Behaviour, vol. 198, Apr. 2023, pp. 59‚Äì72, https://doi.org/10.1016/j.anbehav.2023.01.011. Accessed 2 Mar. 2023. |
| Papio anubis | Mammalia | Direct | dominance | physical contact aggression | 2 | Pritchard, Alexander J., et al. , Individual Differences in Coping Styles and Associations with Social Structure in Wild Baboons (Papio Anubis).‚Äù Animal Behaviour, vol. 198, Apr. 2023, pp. 59,72, doi.org/10.1016/j.anbehav.2023.01.011 |
| Papio cynocephalus | Mammalia | Indirect | group_membership | Number of grooming events between the members of each dyad within a given time frame | 29 | Franz, Mathias, Jeanne Altmann, and Susan C. Alberts. "Knockouts of high-ranking males have limited impact on baboon social networks." Current zoology 61.1 (2015): 107-113. |
| Papio ursinus | Mammalia | Direct | grooming | grooming behaviour (physical contact) | 3 | Roatti, V. et al. (2023), Social network inheritance and differentiation in wild baboons, Royal Society Open Science, 10(5). doi:10.1098/rsos.230219. |
| Philetairus socius | Aves | Indirect | social_projection_bipartite | A network edge was drawn between individuals that used the same nest chambers either for roosting or nest-building at any given time within a series of observations at the same colony in the same year, either together in the nest chamber at the same time or at different times. | 18 | Van Dijk, R., et al. "Cooperative investment in public goods is kin directed in communal nests of social birds." Ecology letters 17.9 (2014): 1141-1148. |
| Saimiri oerstedii | Mammalia | Direct | FE_mating | fluid exchange occurs | 1 | Boinski, S. 1987. Mating Patterns in Squirrel Monkeys (Saimiri oerstedi): Implications for Seasonal Sexual Dimorphism. Behav. Ecol. Sociobiol. 21:13_21. |
| Sapajus apella | Mammalia | Direct | grooming | Based on observation of grooming interaction | 2 | Griffin, Randi H., and Charles L. Nunn. "Community structure and the spread of infectious disease in primate social networks." Evolutionary Ecology 26.4 (2012): 779-800. |
| Sousa sahulensis | Mammalia | Indirect | group_membership | all dolphins sighted within a school were considered associated (Schools were defined as dolphins with relatively close spatial cohesion and involved in similar behavioral activities) | 1 | Hunt, T. N., Allen, S. J., Bejder, L., & Parra, G. J. (2019). Assortative interactions revealed in a fission_fusion society of Australian humpback dolphins. Behavioral Ecology, 30(4), 914-927. |

| | | | | | | |
|---|---|---|---|---|---|---|
| Trachypithecus johnii | Mammalia | Direct | overall_mix | Based on observaions of social interactions including grooming, play behavior and agonistic behavior | 1 | Griffin, Randi H., and Charles L. Nunn. "Community structure and the spread of infectious disease in primate social networks." Evolutionary Ecology 26.4 (2012): 779-800. |
| Trachypithecus johnii | Mammalia | Indirect | group_membership | Based on observaions of social interactions including grooming, play behavior and agonistic behavior | 1 | Griffin, Randi H., and Charles L. Nunn. "Community structure and the spread of infectious disease in primate social networks." Evolutionary Ecology 26.4 (2012): 779-800. |
| Trichosurus cunninghami | Mammalia | Indirect | social_projection_bipartite | The proximity loggers recorded the identity of interacting individuals (based on a threshold proximity set to detect den-sharing events) and the time and length of those interactions. From these data, den-sharing was recorded as a binary variable with _1¬ê representing an instance of day-time den-sharing and _0¬ê representing the use of separate dens for every pairwise combination of individuals on each of the 223 days of data collection. | 1 | Banks, Sam C., et al. "Adaptive responses and disruptive effects: how major wildfire influences kinship-based social interactions in a forest marsupial." Molecular ecology 21.3 (2012): 673-684. |
| Tursiops truncatus | Mammalia | Direct | overall_mix | The overall network does not take behaviour into account but is built from all of the sightings of two dolphins being observed together. | 1 | Gazda, Stefanie, et al. "The importance of delineating networks by activity type in bottlenose dolphins (Tursiops truncatus) in Cedar Key, Florida." Royal Society open science 2.3 (2015): 140263. |
| Tursiops truncatus | Mammalia | Direct | physical_contact | Interactions characterized by repeated incidents of body contact such as rubbing and petting with no consistent direction of movement. | 1 | Gazda, Stefanie, et al. "The importance of delineating networks by activity type in bottlenose dolphins (Tursiops truncatus) in Cedar Key, Florida." Royal Society open science 2.3 (2015): 140263. |
| Tursiops truncatus | Mammalia | Indirect | group_membership | All members of a school were assumed associated. Half-weight index (HWI) was used to quantify the frequency of association among individuals. | 1 | Lusseau, David, et al. "The bottlenose dolphin community of Doubtful Sound features a large proportion of long-lasting associations." Behavioral Ecology and Sociobiology 54.4 (2003): 396-405. |
| Tursiops truncatus | Mammalia | Indirect | group_membership | Interactions characterized by prey capture or persistent incidents of prey searching as indicated by long dives or specialized feeding behaviours with direction shifts between surfacings. | 4 | Gazda, Stefanie, et al. "The importance of delineating networks by activity type in bottlenose dolphins (Tursiops truncatus) in Cedar Key, Florida." Royal Society open science 2.3 (2015): 140263. |
| Zalophus californianus | Mammalia | Direct | group_membership | calculated associations between pairs of individuals as proportion of times they were seen together | 1 | Schakner, Z. A., Petelle, M. B., Tennis, M. J., Van der Leeuw, B. K., Stansell, R. T., & Blumstein, D. T. (2017). Social associations between California sea lions influence the use of a novel foraging ground. Royal Society open science, 4(5), 160820. |

| | | | | | | |
|---|---|---|---|---|---|---|
| Zonotrichia atricapilla | Aves | Indirect | group_membership | A flock was defined as a group of birds within an approximately 5-metre radius. Social networks of flock comembership was constructed where nodes represent individual birds and edges represent the simple ratio association index | 2 | Arnberg, Nina N., et al. "Social network structure in wintering golden-crowned sparrows is not correlated with kinship." Molecular ecology 24.19 (2015): 5034-5044. |
| Zonotrichia atricapilla | Aves | Indirect | group_membership | Flocks were defined as a group of individuals found within a single 5 m radius. For each season, Simple Ratio association index was calculated for each pair of individuals, which ranged from 0 for pairs never seen in the same flock and 1 for pairs always seen in the same flock. | 4 | Shizuka, Daizaburo, et al. "Across-year social stability shapes network structure in wintering migrant sparrows." Ecology letters 17.8 (2014): 998-1007. |

**Table S5:** Sample size for Direct and Indirect interaction type networks.

| Interaction Type | Number of species | Number of Networks |
|---|---|---|
| Direct | 37 | 89 |
| Indirect | 19 | 302 |

**Table S6:** Phylogenetically-controlled Bayesian gaussian model output for strength of relationship disparity of model with formula: *meanlog ~ weighted + clade + (1 | gr(phylo, cov = cov_matrix)) + (1 | species_name)*.

| model term | estimate | std.error | conf.low | conf.high |
|---|---|---|---|---|
| (Intercept) | -0.481 | 0.760 | -1.870 | 1.081 |
| weightedTRUE | -0.010 | 0.526 | -1.081 | 1.011 |
| cladeMammalia | 0.380 | 0.661 | -0.979 | 1.724 |
| sd__(Intercept) phylo | 0.841 | 0.589 | 0.027 | 2.050 |
| sd__(Intercept) species_name | 1.484 | 0.231 | 1.079 | 1.992 |
| sd__Observation Residual | 1.351 | 0.048 | 1.266 | 1.452 |

**Table S7:** Phylogenetically-controlled Bayesian gaussian model output for strength of relationship disparity of model with formula: *meanlog ~ nodes_scale + weighted + (1 | gr(phylo, cov = cov_matrix)) + (1 | species_name)*

| model term | estimate | std.error | conf.low | conf.high |
|---|---|---|---|---|
| (Intercept) | -0.167 | 0.663 | -1.433 | 1.082 |
| nodes_scale | -0.077 | 0.078 | -0.228 | 0.071 |
| weightedTRUE | 0.025 | 0.558 | -1.055 | 1.116 |
| sd__(Intercept) phylo | 0.745 | 0.571 | 0.028 | 2.056 |
| sd__(Intercept) species_name | 1.499 | 0.249 | 1.055 | 2.021 |
| sd__Observation Residual | 1.352 | 0.054 | 1.250 | 1.461 |

**Table S8:** Phylogenetically-controlled Bayesian gaussian model output for strength of relationship disparity of model with formula: *meanlog ~ nodes_scale + weighted + degree_mean_scale + (1 | gr(phylo, cov = cov_matrix)) + (1 | species_name)*

| model term | estimate | std.error | conf.low | conf.high |
|---|---|---|---|---|
| (Intercept) | -0.183 | 0.654 | -1.460 | 1.116 |
| nodes_scale | -0.143 | 0.079 | -0.298 | 0.011 |
| weightedTRUE | -0.150 | 0.575 | -1.278 | 0.964 |
| degree_mean_scale | 0.370 | 0.124 | 0.131 | 0.621 |
| sd__(Intercept) phylo | 0.675 | 0.510 | 0.021 | 1.838 |
| sd__(Intercept) species_name | 1.578 | 0.272 | 1.047 | 2.141 |
| sd__Observation Residual | 1.332 | 0.053 | 1.236 | 1.439 |

**Table S9:** Phylogenetically-controlled Bayesian gaussian model output for strength of relationship disparity of model with formula: *meanlog ~ nodes_scale + weighted + edge_density_scale + (1 | gr(phylo, cov = cov_matrix)) + (1 | species_name)*

| model term | estimate | std.error | conf.low | conf.high |
|---|---|---|---|---|
| (Intercept) | 0.234 | 0.657 | -1.028 | 1.472 |
| nodes_scale | -0.136 | 0.076 | -0.284 | 0.016 |
| weightedTRUE | -0.005 | 0.570 | -1.104 | 1.114 |
| edge_density_scale | -0.568 | 0.121 | -0.796 | -0.330 |
| sd__(Intercept) phylo | 0.625 | 0.493 | 0.017 | 1.815 |
| sd__(Intercept) species_name | 1.701 | 0.249 | 1.230 | 2.209 |
| sd__Observation Residual | 1.301 | 0.051 | 1.206 | 1.403 |

**Table S10:** Phylogenetically-controlled Bayesian gaussian model output for strength of relationship disparity of model with formula: *meanlog ~ nodes_scale + weighted + group_cohesion_scale + (1 | gr(phylo, cov = cov_matrix)) + (1 | species_name)*

| model term | estimate | std.error | conf.low | conf.high |
|---|---|---|---|---|
| (Intercept) | -0.187 | 0.627 | -1.398 | 1.054 |
| nodes_scale | -0.083 | 0.078 | -0.238 | 0.076 |
| weightedTRUE | 0.035 | 0.558 | -1.115 | 1.118 |
| group_cohesion_scale | -0.031 | 0.108 | -0.240 | 0.179 |
| sd__(Intercept) phylo | 0.660 | 0.469 | 0.030 | 1.758 |
| sd__(Intercept) species_name | 1.523 | 0.238 | 1.080 | 2.016 |
| sd__Observation Residual | 1.357 | 0.053 | 1.257 | 1.464 |

**Table S11:** Phylogenetically-controlled Bayesian gaussian model output for strength of relationship disparity of model with formula: *meanlog ~ nodes_scale + weighted + degree_mean_scale + edge_density_scale + group_cohesion_scale + (1 | gr(phylo, cov = cov_matrix)) + (1 | species_name)*

| model term | estimate | std.error | conf.low | conf.high |
|---|---|---|---|---|
| (Intercept) | 0.245 | 0.655 | -1.045 | 1.544 |
| nodes_scale | -0.230 | 0.078 | -0.377 | -0.077 |
| weightedTRUE | -0.236 | 0.588 | -1.360 | 0.924 |
| degree_mean_scale | 0.438 | 0.132 | 0.187 | 0.706 |
| edge_density_scale | -0.884 | 0.153 | -1.187 | -0.577 |
| group_cohesion_scale | 0.360 | 0.134 | 0.092 | 0.625 |
| sd__(Intercept) phylo | 0.623 | 0.545 | 0.017 | 1.928 |
| sd__(Intercept) species_name | 1.834 | 0.285 | 1.324 | 2.431 |
| sd__Observation Residual | 1.251 | 0.050 | 1.161 | 1.357 |

**Table S12:** Phylogenetically-controlled Bayesian gaussian model output for strength of relationship disparity of model with formula: *meanlog ~ nodes_scale + weighted + degree_mean_scale + edge_density_scale + group_cohesion_scale + interaction_type + (1 | gr(phylo, cov = cov_matrix)) + (1 | species_name)*

| model term | estimate | std.error | conf.low | conf.high |
|---|---|---|---|---|
| (Intercept) | 0.380 | 0.684 | -0.974 | 1.704 |
| nodes_scale | -0.230 | 0.076 | -0.376 | -0.084 |
| weightedTRUE | -0.199 | 0.572 | -1.308 | 0.928 |
| degree_mean_scale | 0.441 | 0.137 | 0.180 | 0.716 |
| edge_density_scale | -0.885 | 0.150 | -1.182 | -0.596 |
| group_cohesion_scale | 0.352 | 0.130 | 0.096 | 0.612 |
| interaction_typeDirect | -0.327 | 0.369 | -1.046 | 0.384 |
| sd__(Intercept) phylo | 0.685 | 0.563 | 0.020 | 2.042 |
| sd__(Intercept) species_name | 1.852 | 0.304 | 1.280 | 2.487 |
| sd__Observation Residual | 1.250 | 0.050 | 1.158 | 1.351 |

**Table S13:** Phylogenetically-controlled Bayesian gaussian model output for strength of relationship disparity of model with formula: *meanlog ~ nodes_scale + weighted + degree_mean_scale + edge_density_scale + group_cohesion_scale + interaction_type + clade + (1 | gr(phylo, cov = cov_matrix)) + (1 | species_name)*

| model term | estimate | std.error | conf.low | conf.high |
|---|---|---|---|---|
| (Intercept) | 0.104 | 0.867 | -1.549 | 1.908 |
| nodes_scale | -0.227 | 0.078 | -0.378 | -0.072 |
| weightedTRUE | -0.224 | 0.590 | -1.393 | 0.928 |
| degree_mean_scale | 0.434 | 0.129 | 0.184 | 0.688 |
| edge_density_scale | -0.882 | 0.160 | -1.198 | -0.572 |
| group_cohesion_scale | 0.351 | 0.137 | 0.086 | 0.616 |
| interaction_typeDirect | -0.338 | 0.369 | -1.062 | 0.359 |
| cladeMammalia | 0.385 | 0.730 | -1.065 | 1.794 |
| sd__(Intercept) phylo | 0.845 | 0.691 | 0.023 | 2.580 |
| sd__(Intercept) species_name | 1.795 | 0.308 | 1.206 | 2.441 |
| sd__Observation Residual | 1.252 | 0.049 | 1.160 | 1.350 |

**Table S14:** Phylogenetically-controlled Bayesian gaussian model output for strength of relationship disparity of model with formula: *meanlog ~ nodes_scale + weighted + degree_mean_scale + edge_density_scale + group_cohesion_scale + interaction_type * clade + (1 | gr(phylo, cov = cov_matrix)) + (1 | species_name)*

| model term | estimate | std.error | conf.low | conf.high |
|---|---|---|---|---|
| (Intercept) | 0.039 | 0.893 | -1.648 | 1.905 |
| nodes_scale | -0.232 | 0.075 | -0.381 | -0.089 |
| weightedTRUE | -0.042 | 0.632 | -1.235 | 1.193 |
| degree_mean_scale | 0.455 | 0.141 | 0.196 | 0.752 |
| edge_density_scale | -0.883 | 0.153 | -1.186 | -0.582 |
| group_cohesion_scale | 0.346 | 0.132 | 0.083 | 0.604 |
| interaction_typeDirect | 0.017 | 0.438 | -0.831 | 0.882 |
| cladeMammalia | 0.603 | 0.798 | -1.049 | 2.075 |
| interaction_typeDirect:cladeMammalia | -0.910 | 0.588 | -2.077 | 0.243 |
| sd__(Intercept) phylo | 0.773 | 0.630 | 0.029 | 2.330 |
| sd__(Intercept) species_name | 1.878 | 0.304 | 1.304 | 2.510 |
| sd__Observation Residual | 1.239 | 0.049 | 1.147 | 1.339 |

**Table S15:** Phylogenetically-controlled Bayesian gaussian model output for observed relationship disparity compared to permutation tests, with model formula: *mean_z ~ weighted + (1 | gr(phylo, cov = cov_matrix)) + (1 | species)*

| model term | estimate | std.error | conf.low | conf.high |
|---|---|---|---|---|
| (Intercept) | 0.280 | 0.997 | -1.646 | 2.188 |
| weightedTRUE | -0.866 | 0.842 | -2.480 | 0.887 |
| sd__(Intercept) phylo | 0.780 | 0.692 | 0.017 | 2.660 |
| sd__(Intercept) species_name | 6.999 | 0.698 | 5.783 | 8.487 |
| sd__Observation Residual | 2.027 | 0.077 | 1.881 | 2.183 |

**Table S16:** Phylogenetically-controlled Bayesian gaussian model output for observed relationship disparity compared to permutation tests, with model formula: *mean_z ~ weighted + interaction_type + (1 | gr(phylo, cov = cov_matrix)) + (1 | species)*

| model term | estimate | std.error | conf.low | conf.high |
|---|---|---|---|---|
| (Intercept) | 0.051 | 1.040 | -1.938 | 2.111 |
| weightedTRUE | -0.931 | 0.838 | -2.501 | 0.768 |
| interaction_typeDirect | 0.571 | 0.583 | -0.563 | 1.732 |
| sd__(Intercept) phylo | 0.750 | 0.634 | 0.020 | 2.361 |
| sd__(Intercept) species_name | 7.026 | 0.699 | 5.768 | 8.500 |
| sd__Observation Residual | 2.024 | 0.079 | 1.873 | 2.187 |

**Table S17:** Phylogenetically-controlled Bayesian gaussian model output for observed relationship disparity compared to permutation tests, with model formula: *mean_z ~ weighted + clade + (1 | gr(phylo, cov = cov_matrix)) + (1 | species)*

| model term | estimate | std.error | conf.low | conf.high |
|---|---|---|---|---|
| (Intercept) | 0.422 | 1.217 | -2.058 | 2.804 |
| weightedTRUE | -0.883 | 0.841 | -2.544 | 0.734 |
| cladeMammalia | -0.219 | 0.934 | -2.045 | 1.670 |
| sd__(Intercept) phylo | 0.808 | 0.686 | 0.027 | 2.547 |
| sd__(Intercept) species_name | 6.998 | 0.699 | 5.732 | 8.437 |
| sd__Observation Residual | 2.025 | 0.078 | 1.879 | 2.186 |

**Table S18:** Phylogenetically-controlled Bayesian gaussian model output for observed relationship disparity compared to permutation tests, with model formula: *mean_z ~ weighted + interaction_type + clade + (1 | gr(phylo, cov = cov_matrix)) + (1 | species)*

| model term | estimate | std.error | conf.low | conf.high |
|---|---|---|---|---|
| (Intercept) | 0.210 | 1.232 | -2.287 | 2.633 |
| weightedTRUE | -0.900 | 0.832 | -2.479 | 0.659 |
| interaction_typeDirect | 0.573 | 0.584 | -0.578 | 1.735 |
| cladeMammalia | -0.232 | 0.949 | -2.010 | 1.613 |
| sd__(Intercept) phylo | 0.862 | 0.768 | 0.025 | 2.866 |
| sd__(Intercept) species_name | 7.029 | 0.685 | 5.785 | 8.555 |
| sd__Observation Residual | 2.025 | 0.079 | 1.878 | 2.185 |

**Table S19:** Phylogenetically-controlled Bayesian gaussian model output for observed relationship disparity compared to permutation tests, with model formula: *mean_z ~ weighted + interaction_type * clade + (1 | gr(phylo, cov = cov_matrix)) + (1 | species)*

| model term | estimate | std.error | conf.low | conf.high |
|---|---|---|---|---|
| (Intercept) | 0.141 | 1.276 | -2.372 | 2.678 |
| weightedTRUE | -1.017 | 0.834 | -2.667 | 0.628 |
| interaction_typeDirect | 0.285 | 0.629 | -0.925 | 1.509 |
| cladeMammalia | -0.343 | 0.945 | -2.232 | 1.535 |
| interaction_typeDirect:Mammalia | 1.250 | 0.796 | -0.333 | 2.826 |
| sd__(Intercept) phylo | 0.906 | 0.815 | 0.023 | 3.024 |
| sd__(Intercept) species_name | 7.059 | 0.698 | 5.806 | 8.586 |
| sd__Observation Residual | 2.010 | 0.078 | 1.863 | 2.171 |

**Table S20:** Bayesian gaussian model output for strength of relationship disparity of model with species random effects with the formula: *meanlog ~ 1 + (1 | species_name)*

| model term | estimate | std.error | conf.low | conf.high |
|---|---|---|---|---|
| (Intercept) | -0.252 | 0.252 | -0.740 | 0.238 |
| sd__(Intercept) species_name | 1.581 | 0.225 | 1.184 | 2.065 |
| sd__Observation Residual | 1.354 | 0.052 | 1.256 | 1.462 |

**Table S21:** Bayesian gaussian model output for strength of relationship disparity of model with species random effects with the formula: *meanlog ~ nodes_scale + weighted + (1 | species_name)*

| model term | estimate | std.error | conf.low | conf.high |
|---|---|---|---|---|
| (Intercept) | -0.363 | 0.585 | -1.538 | 0.777 |
| nodes_scale | -0.081 | 0.079 | -0.235 | 0.074 |
| weightedTRUE | 0.108 | 0.587 | -1.035 | 1.259 |
| sd__(Intercept) species_name | 1.601 | 0.222 | 1.198 | 2.067 |
| sd__Observation Residual | 1.353 | 0.051 | 1.259 | 1.457 |

**Table S22:** Bayesian gaussian model output for strength of relationship disparity of model with species random effects with the formula: *meanlog ~ nodes_scale + weighted + degree_mean_scale + (1 | species_name)*

| model term | estimate | std.error | conf.low | conf.high |
|---|---|---|---|---|
| (Intercept) | -0.370 | 0.575 | -1.496 | 0.773 |
| nodes_scale | -0.144 | 0.078 | -0.299 | 0.010 |
| weightedTRUE | -0.100 | 0.584 | -1.248 | 1.062 |
| degree_mean_scale | 0.371 | 0.124 | 0.136 | 0.627 |
| sd__(Intercept) species_name | 1.668 | 0.249 | 1.232 | 2.218 |
| sd__Observation Residual | 1.331 | 0.052 | 1.236 | 1.439 |

**Table S23:** Bayesian gaussian model output for strength of relationship disparity of model with species random effects with the formula: *meanlog ~ nodes_scale + weighted + edge_density_scale + (1 | species_name)*

| model term | estimate | std.error | conf.low | conf.high |
|---|---|---|---|---|
| (Intercept) | 0.072 | 0.598 | -1.101 | 1.238 |
| nodes_scale | -0.137 | 0.078 | -0.288 | 0.017 |
| weightedTRUE | 0.179 | 0.594 | -0.988 | 1.340 |
| edge_density_scale | -0.586 | 0.124 | -0.836 | -0.344 |
| sd__(Intercept) species_name | 1.774 | 0.231 | 1.354 | 2.274 |
| sd__Observation Residual | 1.298 | 0.051 | 1.203 | 1.400 |

**Table S24:** Bayesian gaussian model output for strength of relationship disparity of model with species random effects with the formula: *meanlog ~ nodes_scale + weighted + group_cohesion_scale + (1 | species_name)*

| model term | estimate | std.error | conf.low | conf.high |
|---|---|---|---|---|
| (Intercept) | -0.308 | 0.569 | -1.426 | 0.826 |
| nodes_scale | -0.079 | 0.078 | -0.234 | 0.079 |
| weightedTRUE | 0.107 | 0.565 | -1.001 | 1.271 |
| group_cohesion_scale | -0.038 | 0.106 | -0.249 | 0.165 |
| sd__(Intercept) species_name | 1.611 | 0.215 | 1.217 | 2.069 |
| sd__Observation Residual | 1.354 | 0.052 | 1.259 | 1.459 |

**Table S25:** Bayesian gaussian model output for strength of relationship disparity of model with species random effects with the formula: *meanlog ~ nodes_scale + weighted + degree_mean_scale + edge_density_scale + group_cohesion_scale + (1 | species_name)*

| model term | estimate | std.error | conf.low | conf.high |
|---|---|---|---|---|
| (Intercept) | 0.044 | 0.606 | -1.152 | 1.243 |
| nodes_scale | -0.230 | 0.076 | -0.378 | -0.080 |
| weightedTRUE | -0.019 | 0.584 | -1.148 | 1.114 |
| degree_mean_scale | 0.430 | 0.131 | 0.186 | 0.700 |
| edge_density_scale | -0.903 | 0.157 | -1.205 | -0.597 |
| group_cohesion_scale | 0.363 | 0.132 | 0.113 | 0.627 |
| sd__(Intercept) species_name | 1.901 | 0.260 | 1.446 | 2.456 |
| sd__Observation Residual | 1.251 | 0.048 | 1.162 | 1.350 |

**Table S26:** Bayesian gaussian model output for strength of relationship disparity of model with species random effects with the formula: *meanlog ~ nodes_scale + weighted + degree_mean_scale + edge_density_scale + group_cohesion_scale + interaction_type + (1 | species_name)*

| model term | estimate | std.error | conf.low | conf.high |
|---|---|---|---|---|
| (Intercept) | 0.176 | 0.636 | -1.072 | 1.431 |
| nodes_scale | -0.231 | 0.076 | -0.382 | -0.079 |
| weightedTRUE | 0.039 | 0.601 | -1.140 | 1.234 |
| degree_mean_scale | 0.437 | 0.134 | 0.186 | 0.721 |
| edge_density_scale | -0.891 | 0.158 | -1.195 | -0.579 |
| group_cohesion_scale | 0.352 | 0.133 | 0.089 | 0.614 |
| interaction_typeDirect | -0.318 | 0.359 | -1.042 | 0.390 |
| sd__(Intercept) species_name | 1.923 | 0.280 | 1.423 | 2.530 |
| sd__Observation Residual | 1.248 | 0.050 | 1.154 | 1.349 |

**Table S27:** Bayesian gaussian model output for strength of relationship disparity of model with species random effects with the formula: *meanlog ~ nodes_scale + weighted + degree_mean_scale + edge_density_scale + group_cohesion_scale + clade + (1 | species_name)*

| model term | estimate | std.error | conf.low | conf.high |
|---|---|---|---|---|
| (Intercept) | -0.217 | 0.708 | -1.627 | 1.171 |
| nodes_scale | -0.229 | 0.077 | -0.378 | -0.078 |
| weightedTRUE | -0.112 | 0.603 | -1.279 | 1.079 |
| degree_mean_scale | 0.434 | 0.132 | 0.184 | 0.707 |
| edge_density_scale | -0.903 | 0.155 | -1.203 | -0.607 |
| group_cohesion_scale | 0.361 | 0.131 | 0.113 | 0.620 |
| cladeMammalia | 0.418 | 0.576 | -0.711 | 1.540 |
| sd__(Intercept) species_name | 1.898 | 0.278 | 1.415 | 2.520 |
| sd__Observation Residual | 1.251 | 0.050 | 1.157 | 1.351 |

**Table S28:** Bayesian gaussian model output for strength of relationship disparity of model with species random effects with the formula: *meanlog ~ nodes_scale + weighted + degree_mean_scale + edge_density_scale + group_cohesion_scale + interaction_type + clade + (1 | species_name)*

| model term | estimate | std.error | conf.low | conf.high |
|---|---|---|---|---|
| (Intercept) | -0.075 | 0.740 | -1.547 | 1.370 |
| nodes_scale | -0.231 | 0.076 | -0.384 | -0.083 |
| weightedTRUE | -0.024 | 0.628 | -1.248 | 1.216 |
| degree_mean_scale | 0.432 | 0.140 | 0.172 | 0.719 |
| edge_density_scale | -0.899 | 0.155 | -1.202 | -0.600 |
| group_cohesion_scale | 0.352 | 0.134 | 0.096 | 0.620 |
| interaction_typeDirect | -0.373 | 0.369 | -1.102 | 0.340 |
| cladeMammalia | 0.481 | 0.600 | -0.695 | 1.639 |
| sd__(Intercept) species_name | 1.923 | 0.271 | 1.464 | 2.551 |
| sd__Observation Residual | 1.248 | 0.050 | 1.153 | 1.347 |

**Table S29:** Bayesian gaussian model output for observed relationship disparity compared to permutation tests with species random effects with the formula: *mean_z ~ 1 + (1 | species_name)*

| model term | estimate | std.error | conf.low | conf.high |
|---|---|---|---|---|
| (Intercept) | -0.578 | 0.692 | -1.910 | 0.818 |
| sd__(Intercept) species_name | 6.963 | 0.684 | 5.754 | 8.444 |
| sd__Observation Residual | 2.035 | 0.079 | 1.885 | 2.200 |

**Table S30:** Bayesian gaussian model output for observed relationship disparity compared to permutation tests with species random effects with the formula: *mean_z ~ weighted + interaction_type + (1 | species_name)*

| model term | estimate | std.error | conf.low | conf.high |
|---|---|---|---|---|
| (Intercept) | -0.009 | 1.047 | -2.076 | 2.100 |
| weightedTRUE | -0.915 | 0.881 | -2.619 | 0.847 |
| interaction_typeDirect | 0.600 | 0.572 | -0.518 | 1.701 |
| sd__(Intercept) species_name | 7.039 | 0.699 | 5.801 | 8.537 |
| sd__Observation Residual | 2.027 | 0.081 | 1.878 | 2.195 |

**Table S31:** Bayesian gaussian model output for observed relationship disparity compared to permutation tests with species random effects with the formula: *mean_z ~ weighted + clade + (1 | species_name)*

| model term | estimate | std.error | conf.low | conf.high |
|---|---|---|---|---|
| (Intercept) | 0.471 | 1.214 | -1.882 | 2.746 |
| weightedTRUE | -0.899 | 0.868 | -2.548 | 0.824 |
| cladeMammalia | -0.248 | 0.903 | -1.949 | 1.582 |
| sd__(Intercept) species_name | 7.031 | 0.664 | 5.869 | 8.429 |
| sd__Observation Residual | 2.028 | 0.079 | 1.877 | 2.184 |

**Table S32:** Bayesian gaussian model output for observed relationship disparity compared to permutation tests with species random effects with the formula: *mean_z ~ weighted + interaction_type + clade + (1 | species_name)*

| model term | estimate | std.error | conf.low | conf.high |
|---|---|---|---|---|
| (Intercept) | 0.159 | 1.211 | -2.193 | 2.501 |
| weightedTRUE | -0.901 | 0.880 | -2.599 | 0.819 |
| interaction_typeDirect | 0.553 | 0.577 | -0.576 | 1.688 |
| cladeMammalia | -0.246 | 0.914 | -2.013 | 1.557 |
| sd__(Intercept) species_name | 7.015 | 0.680 | 5.796 | 8.437 |
| sd__Observation Residual | 2.025 | 0.077 | 1.878 | 2.186 |

**Table S33:** Mammalia-only phylogenetically-controlled Bayesian gaussian model output for strength of relationship disparity of model with formula: *meanlog ~ nodes_scale + weighted + degree_mean_scale + edge_density_scale + group_cohesion_scale + (1 | gr(phylo, cov = mammalia_cov_matrix)) + (1 | species_name)*

| model term | estimate | std.error | conf.low | conf.high |
|---|---|---|---|---|
| (Intercept) | 1.188 | 0.774 | -0.387 | 2.715 |
| nodes_scale | 1.972 | 0.206 | 1.564 | 2.370 |
| weightedTRUE | -0.598 | 0.533 | -1.646 | 0.473 |
| degree_mean_scale | -0.398 | 0.173 | -0.713 | -0.041 |
| edge_density_scale | -0.512 | 0.151 | -0.804 | -0.219 |
| group_cohesion_scale | 0.294 | 0.111 | 0.078 | 0.517 |
| sd__(Intercept) phylo | 1.080 | 0.762 | 0.052 | 2.818 |
| sd__(Intercept) species_name | 2.041 | 0.383 | 1.359 | 2.881 |
| sd__Observation Residual | 0.879 | 0.042 | 0.801 | 0.965 |

**Table S34:** Mammalia-only phylogenetically-controlled Bayesian gaussian model output for strength of relationship disparity of model with formula: *meanlog ~ nodes_scale + weighted + degree_mean_scale + edge_density_scale + group_cohesion_scale + interaction_type + (1 | gr(phylo, cov = mammalia_cov_matrix)) + (1 | species_name)*

| model term | estimate | std.error | conf.low | conf.high |
|---|---|---|---|---|
| (Intercept) | 1.402 | 0.769 | -0.028 | 2.830 |
| nodes_scale | 1.938 | 0.205 | 1.519 | 2.307 |
| weightedTRUE | -0.618 | 0.626 | -2.050 | 0.484 |
| degree_mean_scale | -0.331 | 0.170 | -0.674 | 0.005 |
| edge_density_scale | -0.494 | 0.147 | -0.789 | -0.200 |
| group_cohesion_scale | 0.261 | 0.113 | 0.065 | 0.486 |
| interaction_typeDirect | -0.659 | 0.439 | -1.535 | 0.244 |
| sd__(Intercept) phylo | 1.084 | 0.795 | 0.036 | 2.740 |
| sd__(Intercept) species_name | 2.164 | 0.357 | 1.489 | 2.893 |
| sd__Observation Residual | 0.873 | 0.041 | 0.796 | 0.957 |

**Table S35:** Mammalian-only Bayesian gaussian model output for observed relationship disparity compared to permutation tests with species random effects with the formula: *mean_z ~ weighted + (1 | gr(phylo, cov = mammalia_cov_matrix)) + (1 | species_name)*

| model term | estimate | std.error | conf.low | conf.high |
|---|---|---|---|---|
| (Intercept) | 0.680 | 1.105 | -1.492 | 2.869 |
| weightedTRUE | -1.285 | 0.823 | -2.904 | 0.331 |
| sd__(Intercept) phylo | 0.750 | 0.670 | 0.020 | 2.377 |
| sd__(Intercept) species_name | 7.631 | 0.791 | 6.267 | 9.360 |
| sd__Observation Residual | 1.633 | 0.074 | 1.498 | 1.786 |

**Table S36:** Mammalian-only Bayesian gaussian model output for observed relationship disparity compared to permutation tests with species random effects with the formula: *mean_z ~ weighted + interaction_type + (1 | gr(phylo, cov = mammalia_cov_matrix)) + (1 | species_name)*

| model term | estimate | std.error | conf.low | conf.high |
|---|---|---|---|---|
| (Intercept) | 0.102 | 1.089 | -2.101 | 2.212 |
| weightedTRUE | -1.474 | 0.752 | -3.029 | -0.011 |
| interaction_typeDirect | 2.041 | 0.717 | 0.589 | 3.408 |
| sd__(Intercept) phylo | 1.079 | 0.979 | 0.024 | 3.562 |
| sd__(Intercept) species_name | 7.728 | 0.800 | 6.266 | 9.417 |
| sd__Observation Residual | 1.579 | 0.082 | 1.431 | 1.747 |

**Table S37:** Sauropsida-only phylogenetically-controlled Bayesian gaussian model output for strength of relationship disparity of model with formula: *meanlog ~ nodes_scale + weighted + degree_mean_scale + edge_density_scale + group_cohesion_scale + (1 | gr(phylo, cov = sauropsida_cov_matrix)) + (1 | species_name)*

| model term | estimate | std.error | conf.low | conf.high |
|---|---|---|---|---|
| (Intercept) | -0.362 | 0.640 | -1.528 | 0.989 |
| nodes_scale | -0.377 | 0.100 | -0.573 | -0.184 |
| weightedTRUE | -0.466 | 0.703 | -1.806 | 0.945 |
| degree_mean_scale | 1.231 | 0.457 | 0.312 | 2.112 |
| edge_density_scale | -0.720 | 0.317 | -1.344 | -0.096 |
| group_cohesion_scale | 0.407 | 0.361 | -0.299 | 1.094 |
| sd__(Intercept) phylo | 0.603 | 0.488 | 0.023 | 1.859 |
| sd__(Intercept) species_name | 0.960 | 0.478 | 0.121 | 2.016 |
| sd__Observation Residual | 1.573 | 0.119 | 1.370 | 1.818 |

**Table S38:** Sauropsida-only phylogenetically-controlled Bayesian gaussian model output for strength of relationship disparity of model with formula: *meanlog ~ nodes_scale + weighted + degree_mean_scale + edge_density_scale + group_cohesion_scale + interaction_type + (1 | gr(phylo, cov = sauropsida_cov_matrix)) + (1 | species_name)*

| model term | estimate | std.error | conf.low | conf.high |
|---|---|---|---|---|
| (Intercept) | -0.465 | 0.658 | -1.713 | 0.894 |
| nodes_scale | -0.375 | 0.101 | -0.571 | -0.179 |
| weightedTRUE | -0.613 | 0.747 | -2.051 | 0.888 |
| degree_mean_scale | 1.350 | 0.496 | 0.363 | 2.295 |
| edge_density_scale | -0.753 | 0.306 | -1.347 | -0.148 |
| group_cohesion_scale | 0.442 | 0.357 | -0.287 | 1.122 |
| interaction_typeDirect | 0.469 | 0.539 | -0.585 | 1.530 |
| sd__(Intercept) phylo | 0.633 | 0.502 | 0.022 | 1.888 |
| sd__(Intercept) species_name | 0.916 | 0.477 | 0.090 | 1.943 |
| sd__Observation Residual | 1.564 | 0.117 | 1.357 | 1.815 |

**Table S39:** Sauropsida-only Bayesian gaussian model output for observed relationship disparity compared to permutation tests with species random effects with the formula: *mean_z ~ weighted + (1 | gr(phylo, cov = sauropsida_cov_matrix)) + (1 | species_name)*

| model term | estimate | std.error | conf.low | conf.high |
|---|---|---|---|---|
| (Intercept) | -0.333 | 0.912 | -2.139 | 1.468 |
| weightedTRUE | 0.390 | 0.911 | -1.362 | 2.188 |
| sd__(Intercept) phylo | 1.383 | 1.390 | 0.024 | 4.927 |
| sd__(Intercept) species_name | 2.498 | 1.051 | 0.159 | 4.569 |
| sd__Observation Residual | 2.790 | 0.208 | 2.420 | 3.225 |

**Table S40:** Sauropsida-only Bayesian gaussian model output for observed relationship disparity compared to permutation tests with species random effects with the formula: *mean_z ~ weighted + interaction_type + (1 | gr(phylo, cov = sauropsida_cov_matrix)) + (1 | species_name)*

| model term | estimate | std.error | conf.low | conf.high |
|---|---|---|---|---|
| (Intercept) | -0.283 | 0.933 | -2.100 | 1.574 |
| weightedTRUE | 0.395 | 0.902 | -1.384 | 2.077 |
| interaction_typeDirect | -0.043 | 0.750 | -1.492 | 1.392 |
| sd__(Intercept) phylo | 1.225 | 1.231 | 0.027 | 4.480 |
| sd__(Intercept) species_name | 2.617 | 0.983 | 0.433 | 4.599 |
| sd__Observation Residual | 2.793 | 0.200 | 2.435 | 3.221 |

**Table S41:** Variance decomposition for phylogenetically-controlled Bayesian gaussian models of the strength of relationship disparity (natural log transformed as meanlog) of the random effects. Phylogeny denotes the proportion of variance explained by shared evolutionary history, Species identity is the proportion attributable to non-phylogenetic species differences, and within the residual within-species noise. Mean variance extracted from model posterior draws is depicted with 95% credible interval in square brackets. Bayes R2 estimate, estimate error, and 95% credible interval is depicted in the square brackets.

| **Model Formula** | **Phylogeny** | **Species Identity** | **Within** | **Bayes R2** |
|---|---|---|---|---|
| meanlog ~ weighted + clade + (1 \| gr(phylo, cov = cov_matrix)) + (1 \| species_name) | 0.178 [0, 0.544] | 0.45 [0.206, 0.672] | 0.372 [0.228, 0.535] | 0.424 ± 0.032 [0.354, 0.483] |
| meanlog ~ nodes_scale + weighted + (1 \| gr(phylo, cov = cov_matrix)) + (1 \| species_name) | 0.15 [0, 0.549] | 0.469 [0.194, 0.679] | 0.381 [0.224, 0.538] | 0.424 ± 0.033 [0.357, 0.483] |
| meanlog ~ nodes_scale + weighted + degree_mean_scale + (1 \| gr(phylo, cov = cov_matrix)) + (1 \| species_name) | 0.126 [0, 0.487] | 0.508 [0.219, 0.709] | 0.366 [0.223, 0.522] | 0.445 ± 0.033 [0.376, 0.504] |
| meanlog ~ nodes_scale + weighted + edge_density_scale + (1 \| gr(phylo, cov = cov_matrix)) + (1 \| species_name) | 0.106 [0, 0.47] | 0.561 [0.275, 0.738] | 0.333 [0.211, 0.471] | 0.469 ± 0.031 [0.405, 0.524] |
| meanlog ~ nodes_scale + weighted + group_cohesion_scale + (1 \| gr(phylo, cov = cov_matrix)) + (1 \| species_name) | 0.121 [0, 0.454] | 0.488 [0.24, 0.676] | 0.391 [0.249, 0.537] | 0.425 ± 0.032 [0.359, 0.483] |
| meanlog ~ nodes_scale + weighted + degree_mean_scale + edge_density_scale + group_cohesion_scale + (1 \| gr(phylo, cov = cov_matrix)) + (1 \| species_name) | 0.104 [0, 0.493] | 0.608 [0.281, 0.786] | 0.288 [0.173, 0.426] | 0.509 ± 0.030 [0.447, 0.564] |
| meanlog ~ nodes_scale + weighted + degree_mean_scale + edge_density_scale + group_cohesion_scale + interaction_type + (1 \| gr(phylo, cov = cov_matrix)) + (1 \| species_name) | 0.117 [0, 0.529] | 0.603 [0.255, 0.793] | 0.28 [0.166, 0.419] | 0.513 ± 0.029 [0.453, 0.565] |
| meanlog ~ nodes_scale + weighted + degree_mean_scale + edge_density_scale + group_cohesion_scale + interaction_type + clade + (1 \| gr(phylo, cov = cov_matrix)) + (1 \| species_name) | 0.162 [0, 0.652] | 0.563 [0.179, 0.779] | 0.275 [0.141, 0.421] | 0.511 ± 0.029 [0.449, 0.565] |
| meanlog ~ nodes_scale + weighted + degree_mean_scale + edge_density_scale + group_cohesion_scale + interaction_type * clade + (1 \| gr(phylo, cov = cov_matrix)) + (1 \| species_name) | 0.164 [0, 0.635] | 0.577 [0.209, 0.789] | 0.259 [0.138, 0.397] | 0.519 ± 0.029 [0.458, 0.570] |

**Table S42:** Variance decomposition for phylogenetically-controlled Bayesian gaussian models for observed relationship disparity compared to permutation tests for all taxa. Mean variance extracted from model is depicted with 95% credible interval in square brackets. Bayes R2 estimate, estimate error, and 95% credible interval is depicted in the square brackets.

| Model Formula | Phylogeny | Species Identity | Within | Bayes R2 |
|---|---|---|---|---|
| mean_z ~ weighted + (1 \| gr(phylo, cov = cov_matrix)) + (1 \| species_name) | 0.019 [0, 0.119] | 0.903 [0.806, 0.944] | 0.078 [0.051, 0.11] | 0.702 ± 0.017 [0.666, 0.732] |
| mean_z ~ weighted + interaction_type + (1 \| gr(phylo, cov = cov_matrix)) + (1 \| species_name) | 0.018 [0, 0.099] | 0.905 [0.824, 0.945] | 0.077 [0.051, 0.111] | 0.703 ± 0.016 [0.668, 0.732] |
| mean_z ~ weighted + clade + (1 \| gr(phylo, cov = cov_matrix)) + (1 \| species_name) | 0.021 [0, 0.129] | 0.902 [0.794, 0.944] | 0.077 [0.052, 0.111] | 0.702 ± 0.017 [0.665, 0.731] |
| mean_z ~ weighted + interaction_type + clade + (1 \| gr(phylo, cov = cov_matrix)) + (1 \| species_name) | 0.024 [0, 0.139] | 0.9 [0.794, 0.945] | 0.077 [0.05, 0.109] | 0.703 ± 0.017 [0.668, 0.732] |
| mean_z ~ weighted + interaction_type * clade + (1 \| gr(phylo, cov = cov_matrix)) + (1 \| species_name) | 0.026 [0, 0.149] | 0.899 [0.781, 0.945] | 0.075 [0.049, 0.107] | 0.706 ± 0.016 [0.671, 0.735] |

**Table S43:** Variance decomposition for Mammalia-only phylogenetically-controlled Bayesian gaussian models of the strength of relationship disparity (natural log transformed as meanlog) of the random effects. Phylogeny denotes the proportion of variance explained by shared evolutionary history; species identity is the proportion attributable to non-phylogenetic species differences, and within Is the residual within-species noise. Mean variance extracted from model is depicted with 95% credible interval in square brackets. Bayes R2 estimate, estimate error, and 95% credible interval is depicted in the square brackets.

| Model Formula | Phylogeny | Species Identity | Within | Bayes R2 |
|---|---|---|---|---|
| meanlog ~ nodes_scale + weighted + degree_mean_scale + edge_density_scale + group_cohesion_scale + (1 \| gr(phylo, cov = mammalia_cov_matrix)) + (1 \| species_name) | 0.221 [0, 0.708] | 0.656 [0.209, 0.904] | 0.123 [0.061, 0.203] | 0.656 ± 0.025 [0.604, 0.700] |
| meanlog ~ nodes_scale + weighted + degree_mean_scale + edge_density_scale + group_cohesion_scale + interaction_type + (1 \| gr(phylo, cov = mammalia_cov_matrix)) + (1 \| species_name) | 0.205 [0, 0.623] | 0.68 [0.305, 0.906] | 0.115 [0.057, 0.195] | 0.666 ± 0.025 [0.612, 0.710] |

**Table S44:** Variance decomposition for Mammalia-only phylogenetically-controlled Bayesian gaussian models for observed relationship disparity compared to permutation tests. Mean variance extracted from model is depicted with 95% credible interval in square brackets. Bayes R2 estimate, estimate error, and 95% credible interval is depicted in the square brackets.

| Model Formula | Phylogeny | Species Identity | Within | Bayes R2 |
|---|---|---|---|---|
| mean_z ~ weighted + (1 \| gr(phylo, cov = mammalia_cov_matrix)) + (1 \| species_name) | 0.016 [0, 0.09] | 0.94 [0.866, 0.969] | 0.044 [0.028, 0.065] | 0.811 ± 0.011 [0.787, 0.831] |
| mean_z ~ weighted + interaction_type + (1 \| gr(phylo, cov = mammalia_cov_matrix)) + (1 \| species_name) | 0.031 [0, 0.162] | 0.929 [0.808, 0.971] | 0.04 [0.025, 0.061] | 0.821 ± 0.012 [0.797, 0.841] |

**Table S45:** Variance decomposition for Sauropsida-only phylogenetically-controlled Bayesian gaussian models of the strength of relationship disparity (natural log transformed as meanlog) of the random effects. Phylogeny denotes the proportion of variance explained by shared evolutionary history; species identity is the proportion attributable to non-phylogenetic species differences, and within Is the residual within-species noise. Mean variance extracted from model is depicted with 95% credible interval in square brackets. Bayes R2 estimate, estimate error, and 95% credible interval is depicted in the square brackets.

| Model Formula | Phylogeny | Species Identity | Within | Bayes R2 |
|---|---|---|---|---|
| meanlog ~ nodes_scale + weighted + degree_mean_scale + edge_density_scale + group_cohesion_scale + (1 \| gr(phylo, cov = sauropsida_cov_matrix)) + (1 \| species_name) | 0.122 [0, 0.528] | 0.247 [0.004, 0.606] | 0.631 [0.309, 0.906] | 0.429 ± 0.058 [0.302, 0.526] |
| meanlog ~ nodes_scale + weighted + degree_mean_scale + edge_density_scale + group_cohesion_scale + interaction_type + (1 \| gr(phylo, cov = sauropsida_cov_matrix)) + (1 \| species_name) | 0.134 [0, 0.548] | 0.234 [0.002, 0.59] | 0.632 [0.306, 0.898] | 0.433 ± 0.056 [0.314, 0.530] |

**Table S46:** Variance decomposition for Sauropsida-only phylogenetically-controlled Bayesian gaussian models for observed relationship disparity compared to permutation tests. Mean variance extracted from model is depicted with 95% credible interval in square brackets. Bayes R2 estimate, estimate error, and 95% credible interval is depicted in the square brackets.

| Model Formula | Phylogeny | Species Identity | Within | Bayes R2 |
|---|---|---|---|---|
| mean_z ~ weighted + (1 \| gr(phylo, cov = sauropsida_cov_matrix)) + (1 \| species_name) | 0.159 [0, 0.751] | 0.387 [0.001, 0.723] | 0.453 [0.198, 0.745] | 0.382 ± 0.065 [0.243, 0.495] |
| mean_z ~ weighted + interaction_type + (1 \| gr(phylo, cov = sauropsida_cov_matrix)) + (1 \| species_name) | 0.132 [0, 0.695] | 0.410 [0.007, 0.726] | 0.458 [0.219, 0.743] | 0.384 ± 0.065 [0.245, 0.498] |
